# ALDH1 subtype-specific inhibitor targets key cancerous epithelial cell populations in aggressive subtypes of breast cancer

**DOI:** 10.1101/2024.10.18.619128

**Authors:** Raquel Pequerul, Andrada Constantinescu, Bassam Janji, Akinchan Kumar, Xavier Parés, Oscar Palacios, Delphine Colignon, Axelle Berrou, Guy Fournet, Paul Berchard, Guillaume Martin, Ismail Ceylan, Rocio Rebollido-Ríos, Jaume Farrés, Mileidys Perez-Alea

## Abstract

Aggressive breast cancer subtypes, particularly triple-negative breast cancer (TNBC), lack effective targeted therapies, requiring novel approaches. This study focuses on the aldehyde dehydrogenase 1A (ALDH1A) subfamily, comprising ALDH1A1, ALDH1A2, and ALDH1A3, and their roles in tumor biology and the tumor microenvironment. Comprehensive transcriptomic and single-cell analyses revealed subtype- and cell-specific expression patterns of ALDH1A isoforms, with *ALDH1A3* predominantly expressed in epithelial cancer cells of basal-like tumors, while *ALDH1A1* and *ALDH1A2* were expressed in stromal and immune-associated subpopulations. Guided by these findings, we developed ABD0171, a selective ALDH1A3 inhibitor that demonstrated potent isoform-specific activity. ABD0171 effectively disrupted key pathways in TNBC cells *in vitro*, including IL6/JAK/STAT3, tPA and Src/FAK, and exhibited superior selectivity, robust antitumor and antimetastatic effects, and a favorable safety profile *in vivo*. These results establish ALDH1A3 as a promising therapeutic target and validate ABD0171 as a candidate to address current challenges in aggressive breast cancers.

## INTRODUCTION

Breast cancer is the most prevalent malignancy among women worldwide, with metastatic progression remaining the leading cause of mortality^1^. This disease exhibits extensive clinical and molecular heterogeneity, which complicates treatment strategies. Over recent years, the classification of breast cancer has been expanded beyond the four major subtypes defined by the presence or absence of estrogen receptor (ER), progesterone receptor (PR), and human epidermal growth factor receptor 2 (HER2)^2–5^. Triple-negative breast cancer (TNBC), defined by the lack of expression of ER, PR and HER2, represents a particularly challenging subset^6^. Advanced molecular profiling has further subdivided breast cancers into intrinsic subtypes, including Luminal A, Luminal B, HER2-enriched, Basal-like, and normal-like entities, providing greater insight into tumor biology and therapeutic opportunities^4,7,8^. Notably, TNBCs, which frequently align with the Basal-like subtype, account for ∼15% of invasive cases. These tumors are characterized by high aggressiveness, a lack of targeted therapies, and poor clinical outcomes, making them a critical focus for research and therapeutic development^6,9^.

The aldehyde dehydrogenase 1A (ALDH1A) subfamily of enzymes-comprising ALDH1A1, ALDH1A2, and ALDH1A3-has gained significant attention as a key player in breast cancer biology, particularly in TNBC. Historically, ALDH1 activity has served as a biomarker of cancer stem cells, and has been functionally implicated in tumor initiation, progression, and therapy resistance, primarily through its roles in oxidative stress mitigation and detoxification of reactive aldehydes derived from aberrant metabolism or cytotoxic agents. Recent findings, however, underscore isoform-specific contributions: ALDH1A1 expression is linked to chemoresistant breast cancer stem cells^10–12^, ALDH1A2 is proposed as a diagnostic biomarker for early breast cancer^13^, and ALDH1A3 is strongly associated with proliferation and poor prognosis in triple negative breast cancer (TNBC)^11,14–18;^ likely due to their role in retinoic acid (RA) synthesis—a crucial regulator of differentiation and proliferation^19^. Despite mounting evidence supporting ALDH1A isoforms as important biomarkers and therapeutic targets in breast cancer, the precise contribution of each isoform to tumor pathogenesis, including their distribution within the tumor microenvironment, and their expression patterns across breast cancer molecular subtypes remain poorly understood. This gap in knowledge has hindered the development of isoform-selective therapies to effectively target hard-to-treat breast cancer subtypes like TNBC.

In this study, we investigated the distribution and abundance of *ALDH1A1*, *ALDH1A2*, and *ALDH1A3* across breast cancer molecular subtypes and at single-cell resolution within the tumor microenvironment. Guided by these insights, we developed a series of ALDH1 inhibitors based on the structure of DIMATE (4-dimethylamino-4-methyl-pent-2-ynthioic acid-S-methyl ester), an ALDH inhibitor currently in Phase I clinical trial (“ODYSSEY”, NCT05601726) for acute myeloid leukemia, with demonstrated preclinical efficacy in leukemia, melanoma, and non-small cell lung cancer. The lead compound, ABD0171, demonstrated potent isoform-specific activity, disrupted key pathways in TNBC *cells in vitro*, and exhibited superior selectivity, robust antitumor and antimetastatic effects, and a favorable safety profile *in vivo*.

## RESULTS

### ALDH1A isoforms exhibit specific expression patterns across molecular subtypes and cell populations in breast tumors

We preprocessed and integrated three independent RNA-seq datasets from The Cancer Genome Atlas (TCGA)^20,21^ to generate a comprehensive bulk RNA-seq dataset comprising 1103 primary human breast invasive carcinoma samples (Fig.1a). Unsupervised clustering of the integrated dataset, incorporating 59 genes-including the PAM50 signature,^22,23^ claudin-low associated-genes^24^ and *ALDH1A1*, *ALDH1A2*, and *ALDH1A3* classified the tumors into four intrinsic molecular subtypes according to the PAM50 classifier: Basal-like (17.8%), HER2-enriched (36.4%), Luminal A (25.4%) and Luminal B (18.5%). Additionally, a small subset (1.9 %) exhibited claudin-low molecular features, consistent with the reported prevalence of claudin-low tumors, which account for 1.5% to 14% of all breast tumors (Fig. 1b, Supplementary Data 1A).

**Figure 1.**
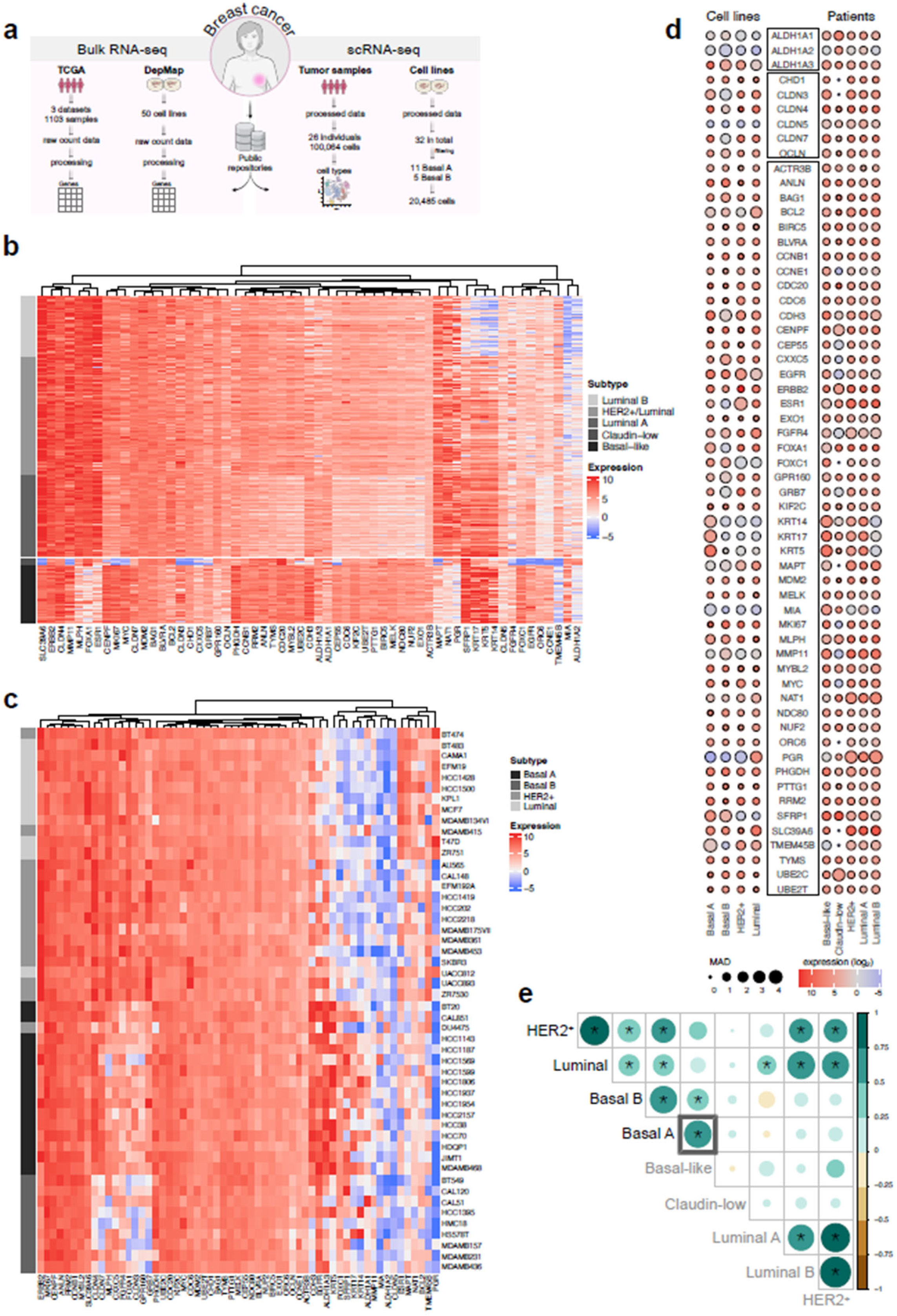
Data-based transcriptomic profiling of breast cancer cell lines and patient tumors. **(a)** Overview of data collection, detailing the number of human cell lines and patient samples analyzed. **(b-c)** Heatmaps showing the differential expression of PAM50 genes and ALDH1 isoforms (*ALDH1A1*, *ALDH1A2*, and *ALDH1A3*) in cell lines and patient samples, respectively, with molecular subtypes represented as annotated rows to the left of the heatmaps. **(d)** Median bulk expression of 59 genes in breast cancer cell lines and patient samples, grouped by molecular classification. Dot size represents median absolute deviation (MAD), with color intensity reflecting log_2_ expression levels. **(e)** Spearman correlation of gene expression between groups in **(d)**, with significant correlation (p-value ≤ 0.01) marked by stars. A gray square highlights a strong positive correlation (ρ = 0.71) between Basal A (cell lines) and Basal-like (patient samples) subtypes.

We also retrieved RNA-seq data for 50 human breast cancer cell lines from the Dependency Map (DepMap)^25^ portal and performed the subsequent preprocessing steps. Unsupervised clustering of these cell lines revealed four distinct molecular subgroups, aligning with established classifications of breast cell lines ^26^: Basal A (32%), Basal B (18%), HER2-enriched (26%), and Luminal (24%) (Fig. 1a, c, Supplementary Data 1B). We used these classifications to analyze the expression of *ALDH1A1*, *ALDH1A2* and *ALDH1A3* across breast cancer subtypes, aiming to clarify their expression profile and assess how well cell lines reflect the primary tumors. Among the cell lines, *ALDH1A3* was the most predominantly expressed isoform, with significantly higher expression in Basal A compared to Basal B, HER2-enriched and luminal subtypes (adjusted *p*-value < 0.001, Fig.1d and Supplementary Data 1C). This finding was consistent with ALDH1A3 protein expression profiles (Supplementary Fig. 1a) and supported by previous studies that associate high ALDH1A3 expression with TNBC^16,17^, an entity identified by surrogate immunohistochemistry as negative for hormone receptors and HER2, reported to overlap significantly (75%) with the Basal-like subtype^27^. Across all breast cancer cell lines, *ALDH1A1* and *ALDH1A2* were expressed at low levels, regardless of molecular subtype. Basal A cell lines have been reported to closely resemble the Basal-like tumors in clinical samples, sharing key characteristics such as epithelial properties and highly proliferative capacity^28^. We observed a strong positive correlation (ρ = 0.71) between Basal A subtype in cell lines and the Basal-like subtype in patient samples, further confirming their shared molecular features (Fig.1e). Notably, this similarity extends to the significant expression of *ALDH1A3* in both Basal A cell lines and Basal-like breast cancer patients (adjusted *p*-value < 0.001, Fig. 1d and Supplementary Data 1D and 1E). *ALDH1A2* expression remained low across all intrinsic molecular subtypes in patient samples, reflecting a trend like that observed in the cell line data. In contrast, *ALDH1A1* exhibited higher expression in all patient subtypes, suggesting that its expression may originate from diverse cell types within the tumor microenvironment, which are not fully represented in traditional cell line models (Fig. 1d).

Interestingly, the specific subset of patients with claudin-low tumors exhibited significant expressions of both *ALDH1A1* and *ALDH1A2* compared to the other tumor types (all adjusted *p*-values < 0.001, Fig. 1d and Supplementary Data 1E). Although claudin-low tumors are challenging to model *in vitro* due to their characteristic stromal compartment, three cell lines (HCC1143, MDAMB436 and DU4475) that had been previously identified as molecularly and phenotypically similar to claudin-low tumors^5^ also displayed high ALDH1A2 protein levels and double positivity for ALDH1A1 protein (Supplementary Fig. 1a). Together, these observations suggest that claudin-low breast cancers possess a distinct ALDH1A expression profile.

To dissect the individual contributions of different cell types to the overall *ALDH1A* gene expression in both cell lines and tumor populations, we analyzed single-cell RNA sequencing (scRNA-seq) data collected from two publicly independent studies targeting breast cancer cell lines and patients, respectively. One study was based on 32 breast cancer cell lines^29^ while another based on primary tumor biopsies from 26 patients with invasive breast carcinoma including a total of 100,064 cells^30^. The earlier study, which included the breast cancer cell lines, revealed that basal phenotype cell lines form a distinct cluster, separate from the luminal and HER2-enriched groups^29^. Focusing only on the 16 basal cell lines including a total of 20,485 cells (Fig. 2a and Supplementary Data 1F), we identified a distinct cluster of Basal B cells, with two Basal A cell lines (HCC1143 and JIMT1) mixed within a broader Basal A population (Fig. 2b). Differential gene expression analysis between these basal phenotypes confirmed that *ALDH1A3* is highly expressed in the Basal A subtype (Fig. 2b), consistent with the findings shown in Fig. 1d. The expression levels of *ALDH1A1* and *ALDH1A2* remain very low or nearly absent in both Basal A and Basal B cell lines subtypes, also aligning with Fig. 1d.

**Figure 2.**
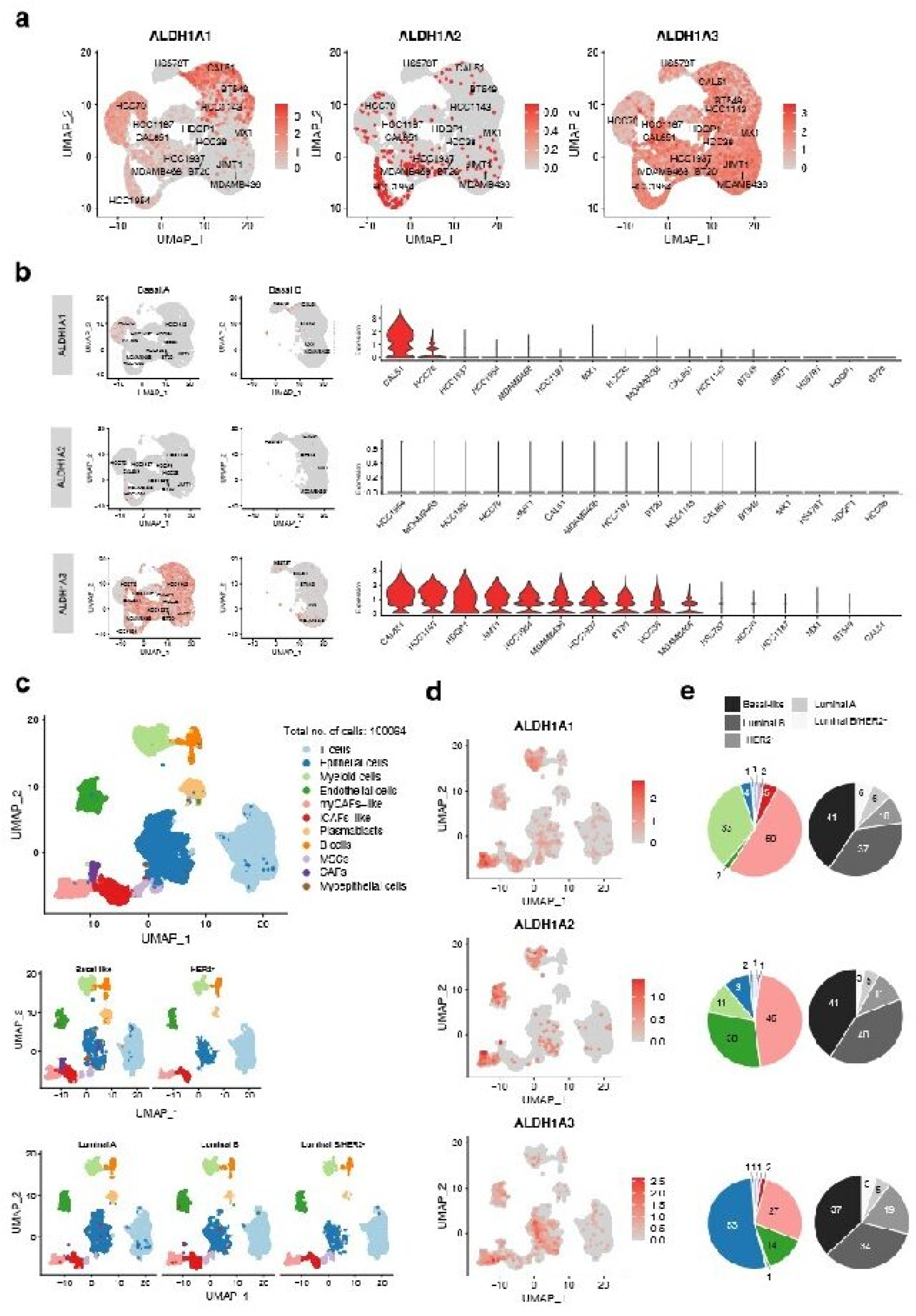
Single-cell RNA sequencing reveals distinct ALDH1A isoform expression across tumor populations in breast cancer. **(a,b)** Expression patterns of *ALDH1A1, ALDH1A2*, and *ALDH1A3* across 20,485 cells from 16 Basal-like classified breast cancer cell lines, visualized using feature and violin plots. Classification into Basal A and B subgroups highlights intra-group heterogeneity and variability. **(c)** Single-cell analysis of 26 patient samples encompassing 100,064 cells, detailing the *ALDH1A* expression landscape. **(d)** Distribution of cell populations by molecular classification, illustrating cellular heterogeneity within tumors. **(e)** Feature plots display gene expression distribution and intensity for *ALDH1A1*, *ALDH1A2*, and *ALDH1A3* among individual cells (left). Corresponding pie charts depict comparative gene expression of each ALDH1A isoform, indicating the percentage of cells grouped by population and molecular subtype (right). Cell types analyzed include cancer-associated fibroblasts (CAFs), myofibroblasts (myCAFs), inflammatory CAFs (iCAFs), and mesenchymal stem cells (MSCs).

Next, we explored the heterogeneity of the ALDH1A subfamily across breast cancer subtypes by analyzing the patient samples^30^. Using canonical cell markers, we identified 10 major cell types along with a small population of myoepithelial cells (Fig. 2c). The major cell types were annotated as epithelial, stromal (including endothelial, mesenchymal stem cells (MSCs), cancer-associated fibroblasts (CAFs), myofibroblastic CAFs-like (my-CAFs-like) and inflammatory CAF-like (iCAFs-like) and immune cells (T-cells, myeloid cells, B cells and plasmablasts). These cell populations were found across all five intrinsic molecular subtypes, except for CAFs and myoepithelial cells, which were exclusively detected in the Basal-like subtype (Fig. 2c and Supplementary Data 1G).

Our analyses revealed unique expression patterns of the *ALDH1A* isoforms within specific cell populations (Fig. 2d, e). Among the three *ALDH1A* isoforms, only *ALDH1A1* was differentially expressed in the myCAFs-like population. *ALDH1A1* was expressed in 2,853 cells, *ALDH1A3* in 1,956 cells, and *ALDH1A2* in only 502 cells. Comparative analysis of isoform expression patterns across cell populations and molecular subtypes is visualized in two sets of pie charts in Fig. 2e. The data reveal unique expression profiles for *ALDH1A1, ALDH1A2* and *ALDH1A3* across different cell types within breast tumors. Notably, *ALDH1A1* and *ALDH1A3* are uniquely expressed in myeloid cells (33% and 1%, respectively) and epithelial cells (4% and 53%, respectively). *ALDH1A1* and *ALDH1A2* display similar positivity within the myCAFs-like population (50% and 46%, respectively) while *ALDH1A*3 is 27%. Approximately, 70-80% of the cells expressing *ALDH1A1, ALDH1A2*, or *ALDH1A3* originate from Basal-like and Luminal B subtypes. Interestingly, the HER2-enriched subtype contributes to 19% of the ALDH1A3-expressing cells, a higher proportion compared to its representation in ALDH1A1 and ALDH1A2 positive expression profiles.

Overall, these insights emphasize the complex, cell population-specific expression patterns of ALDH1A isoforms in breast tumors, underscoring their unique roles in different breast cancer subtypes and their potential impact on targeted therapies.

### Novel compounds with selective inhibitory activity towards ALDH1A3 show enhanced cytotoxicity in basal phenotype breast cancer

Building upon our characterization of the ALDH1A landscape in breast tumors, we next sought to explore the therapeutic potential of targeting these enzymes. To this end, we designed and synthesized a series of novel compounds, categorized into different families by their chemical substituents (Fig. 3a). DIMATE and MATE represent prior α,β-acetylenic aminothiolester compounds featuring a pentynethioic acid backbone^31^. In previous efforts to generate potent ALDH inhibitors, we explored pharmacological modulation of DIMATE at the N atom with alkyl, aryl or heterocyclic substituents (series I and II) and/or the substitution of the sulphur atom of the thiolester functional group with oxygen (series III). However, these approaches did not yield increased potency or selectivity for this family of compounds^31^. Here, we performed novel targeted structural modifications mainly at the nitrogen atom of the original aminothiolester core structure. The new compounds were designed with a -N-C-C-O-pattern which corresponds to the morpholine ring opening in MATE (IV, V, VI, and VII series) (Fig. 3a)^32,33^. We next evaluated their effects on the enzymatic activity of recombinant human ALDH1A1, ALDH1A2, and ALDH1A3, as well as ALDH2 (the isoform most closely related to ALDH1A and known for its cardioprotective effects in the heart) and the critical corneal ALDH3A1, through a single-point enzymatic inhibition analysis (Fig. 3b and Supplementary Data 2A). The most effective compounds against ALDH1A1, ALDH1A2 and ALDH1A3 activity belonged to series V of analogs, characterized by a (arylmethoxy)ethylamino substituents compared to series IV (alkylmethoxy)ethylamino substituents (e.g. ABD0118). Notably, three compounds within the same family V, containing an (heteroarylmethoxy)ethylamino group substitution instead (i.e. ABD0257, ABD0258 and ABD0260), were as ineffective as, for example, ABD0118 in inhibiting ALDH1A activity. This suggests that the aryl moiety is a crucial modification for achieving enhanced and selective inhibitory activity against all three human ALDH1A isoforms, while preserving the functions of ALDH2 and ALDH3A1.

**Figure 3.**
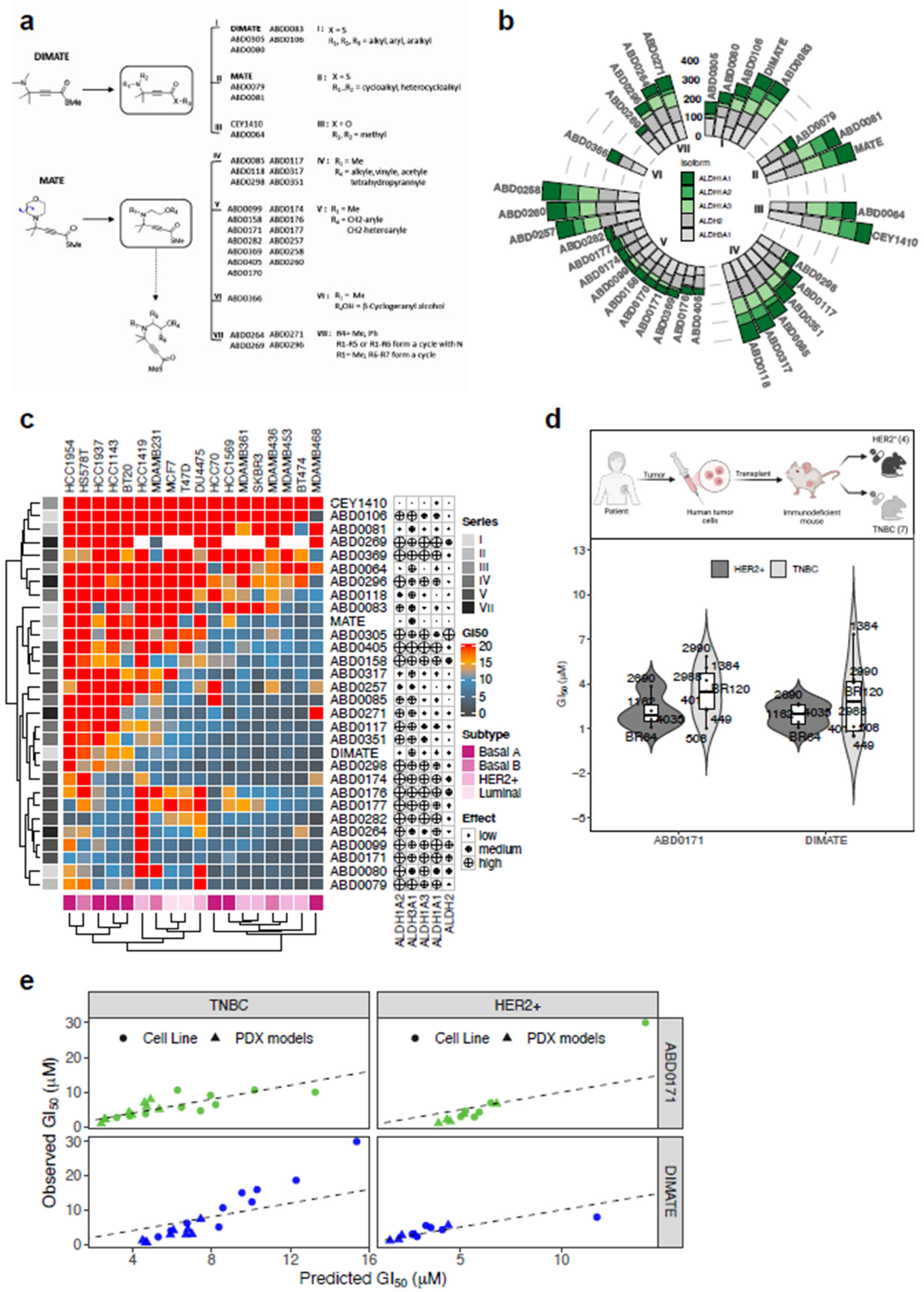
Novel chemical compounds targeting ALDH1A isoforms and their cytotoxic effects on breast cancer cell lines and patient-derived xenograft (PDX) models. **(a)** Schematic representation of the structural analogs of DIMATE and MATE developed for selective ALDH1A inhibition. **(b)** Circular stacked barplots showing the remaining activity (%) of compounds against five aldehyde dehydrogenase isoforms. ALDH1A1, ALDH1A2, and ALDH1A3 are represented in shades of green (dark to light), while ALDH2 and ALDH3A1 are depicted in shades of gray. The segment lengths indicate the proportion of remaining activity, providing a comparative view of compound efficacy. **(c)** Heatmap of cytotoxic activity (GI50 values) of the compounds, grouped by series number and their inhibitory effects on ALDH1 isoforms, evaluated across 18 breast cancer cell lines stratified by molecular subtypes. **(d)** Experimental design for 3D clonogenic assays using patient-derived cells from HER2-enriched and TNBC xenograft models (top). Violin plots display the GI50 distribution for DIMATE and ABD0171 in these models (bottom). **(e)** Linear mixed-effects model analysis showing predicted versus observed cytotoxic effects (GI50 values) for DIMATE and ABD0171 in TNBC and HER2- enriched cell lines and PDX models.

To investigate the anticancer potential of synthetized compounds, we conducted cytotoxic drug-response assays using a panel of 18 human breast cancer cell lines representing the four major molecular subtypes (Supplementary Data 3A). We reported the results as half-growth inhibition (GI_50_) values calculated from the corresponding dose-response curves (Supplementary Fig. 1b, Supplementary Data 3B) and represented them as a heatmap to highlight compounds with enhanced activity against different subtypes alongside their ALDH inhibitory profiles (Fig. 3c). Compounds in series V, characterized by aryl substitutions-except for compound ABD0369, which features a bicyclic aromatic hydrocarbon (i.e. naphthalene)-demonstrated the most potent cytotoxic effects. ABD0171 exhibited exceptional potency, with GI_50_ values consistently below 10 µM in 95% of tested cell lines (Supplementary Data 3B).

Additionally, compounds ABD0080, featuring a benzyl substitution on nitrogen, and ABD0079, forming a heterocycle with nitrogen, also showed notable efficacy. Importantly, these two compounds, along with most aryl-substituted compounds in series V (except ABD0405 and ABD0158) exhibited selective cytotoxic activity toward Basal A and Basal B cell lines, with ABD0171 being the most effective.

In contrast, compounds from series I, II, III and IV, particularly those with minimal inhibitory effects on ALDH1A1 and ALDH1A3, showed little to no cytotoxic activity against the majority of basal phenotype cell lines. These compounds were notably ineffective against BT20, HCC1143, HCC1937, and HCC1954 Basal A cell lines, as well as the HER2-enriched HCC1419 cells, which express high ALDH1A3 protein levels. This suggests a specific association between the compounds’ efficacy and the ALDH1A3 inhibition. Intriguingly, HER2-enriched cells, which co-expressed ALDH1A1 and ALDH1A3 in 4 out of 5 cell lines, showed high sensitivity to DIMATE and compounds within family IV, despite their weaker ALDH1A inhibitory effects. This finding raises the possibility that the single-point enzymatic analysis may not fully capture their inhibitory effects on ALDH1A1 and/or ALDH1A3, or that their cytotoxicity may result from inhibition of another ALDH isoform or an off-target mechanism critical for HER2-enriched cell survival.

Next, as shown in Fig. 3d, we evaluated the impact of ABD0171 and DIMATE on the self-renewal and sustain proliferation of breast cancer patient-derived xenograft (PDX) cells. These PDX models were characterized by immunochemistry as HER2-enriched subtype (n=4) and triple-negative breast cancer (TNBC, n=7). Both compounds displayed similar inhibitory effects on long-term proliferation in colony formation assays across both PDX subtypes, suggesting minimal differences in their tumorigenic potential in this context (Supplementary Data 3C).

To further investigate the relationship between predicted and observed drug response in both PDX models and cancer cell lines, we employed a linear mixed-effects model (LMM). This analysis incorporated GI_50_ values from TNBC and HER2-enriched PDX as well as cell lines. The average GI_50_ for ABD0171 in Basal A cell lines was used as the baseline for the LMM analysis, given their demonstrated sensitivity to the compound. As shown in Fig. 3e, the LMM revealed a robust relationship between predicted and observed GI_50_ values for both TNBC and HER2-enriched models treated with ABD0171 and DIMATE. In TNBC models, ABD0171 showed a strong predictive performance, as evidenced by the close alignment of observed and predicted GI_50_ values for most cell lines and PDX models (Fig. 3e, Supplementary Data 3D). In contrast, DIMATE-treated TNBC models showed greater variability, indicating that while potent, its effects might be modulated by additional factors not fully captured by the model. In HER2-enriched models, both compounds performed similarly. DIMATE exhibited the strongest agreement between predicted and observed GI_50_ values, whereas ABD0171, though effective, showed a few outliers in PDX models, suggesting the presence resistant mechanisms or differential drug activity in HER-2-enriched cells that warrants further mechanistic investigation.

### ABD0171 is a potent irreversible inhibitor and specific affinity-label against ALDH1A3

Given the strong ALDH1A inhibitory effect and potent cytotoxicity of ABD0171 against Basal-like cancer cells, we conducted a detailed enzymatic and biochemical characterization of ABD0171 alongside DIMATE. We determined the IC50 values and the type of inhibition for both compounds against ALDH1A1, ALDH1A2, ALDH1A3, ALDH2 and ALDH3A1 isoforms, using both routine (hexanal) and physiological (all-*trans*-retinaldehyde, atRAL) substrates and for ALDH3A1 its preferred 4-nitrobenzaldehyde substrate (Table 1 and Supplementary Table S1). These assays revealed that inhibition was contingent on the order of reagent addition and incubation times. No inhibition could be observed when the cofactor was added prior to the inhibitors, suggesting the existence of steric or conformational impediments. Both compounds exhibited time-dependent inhibition of all ALDH isoforms (Fig. 4a and Supplementary Fig. 2a), except for ALDH3A1 (Supplementary Fig. 2a), suggesting that DIMATE and ABD0171 act as irreversible inhibitors for all the other isoforms. The ALDH1A IC50 values with the physiological atRAL substrate ranged from 0.34 to 6 µM for ABD0171 and from 5.7 to 25 for DIMATE, after a 20-min preincubation (Table 1). In contrast, IC50 values for ALDH2 and ALDH3A1 remained substantially higher for both inhibitors, highlighting their selectivity against ALDH1A isoforms (Supplementary Table S1). Additionally, kinetic analysis confirmed non-competitive inhibition of ALDH3A1 by both DIMATE and ABD0171 with *K_i_*values of 253 ± 48 µM and 233 ± 16 µM, respectively (Supplementary Fig. 2a and Supplementary Table S1).

**Figure 4.**
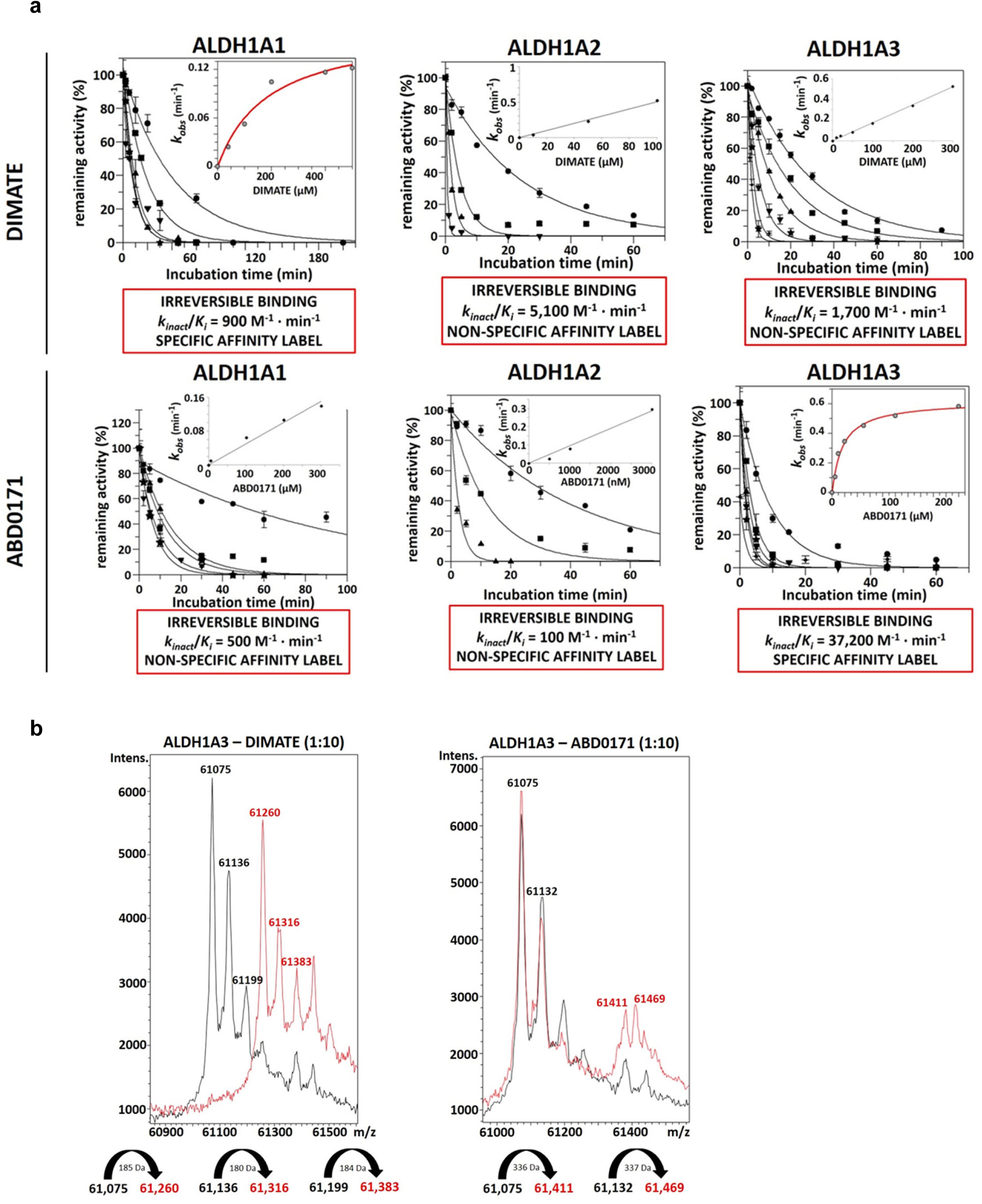
Kinetic analyses and mass spectrometry validation of DIMATE and ABD0171 as ALDH1A inhibitors. **(a)** *k_inact_/K_i_* values calculated from the plots of *k_obs_ versus* concentrations of DIMATE and ABD0171 for ALDH1A1, ALDH1A2 and ALDH1A3 (inner plots). For DIMATE, tested concentration ranges were 0–500 µM for ALDH1A1, 0–200 µM for ALDH1A2, and 0–300 µM for ALDH1A3. For ABD0171, concentration ranges were 0–300 µM for ALDH1A1, 0–3 µM for ALDH1A2, and 0–200 µM for ALDH1A3. Enzymatic activities were measured at saturating substrate concentrations (250 µM hexanal) in the corresponding reaction buffer. Data are presented as mean ± standard deviation. **(b)** ESI-MS spectra of ALDH1A3 treated with DIMATE (left) and ABD0171 (right). Single-charged peaks (1+) were observed at *m/z* 61075–61119. Red signals represent ALDH1A3 treated with 167 µM DIMATE or ABD0171 (at an enzyme concentration of 16.7 µM), while black signals correspond to the untreated ALDH1A3 control. The spectra confirm irreversible binding of ALDH1A3 to DIMATE and ABD0171. Samples were prepared in 100 mM ammonium bicarbonate (pH 7.5) and pre-incubated for 45 min at 37°C.

**TABLE 1.** IC_50_ (µM) and *k_inact_*/*K_i_* (M^−1^ · min^−1^) values of DIMATE and ABD0171 for ALDH1A1, ALDH1A2 and ALDH1A3 using hexanal and all-*trans*-retinaldehyde as substrates.

| COMPOUND | ALDH1A1 |  |  | ALDH1A2 |  |  | ALDH1A3 |  |  |  |
| --- | --- | --- | --- | --- | --- | --- | --- | --- | --- | --- |
|  | atRAL | HEXANAL |  | atRAL | HEXANAL |  | atRAL |  | HEXANAL |  |
| | $IC_{50}$<br>($\mu M$) | $IC_{50}$<br>($\mu M$) | $k_{inact}/K_i$<br>($M^{-1} \cdot min^{-1}$) | $IC_{50}$<br>($\mu M$) | $IC_{50}$<br>($\mu M$) | $k_{inact}/K_i$<br>($M^{-1} \cdot min^{-1}$) | $IC_{50}$<br>($\mu M$) | $k_{inact}/K_i$<br>($M^{-1} \cdot min^{-1}$) | $IC_{50}$<br>($\mu M$) | $k_{inact}/K_i$<br>($M^{-1} \cdot min^{-1}$) |
| **DIMATE** | $25 \pm 2$ | $37 \pm 5$ | 900 | $6.4 \pm 0.9$ | $18 \pm 4$ | 5,100 | $5.7 \pm 0.5$ | 80,100 | $20 \pm 2$ | 1,700 |
| **ABD0171** | $6.6 \pm 1$ | $3.8 \pm 1.1$ | 500 | $0.34 \pm 0.03$ | $0.57 \pm 0.1$ | 100 | $0.79 \pm 0.2$ | 115,400 | $3.4 \pm 0.1$ | 37,200 |

$IC_{50}$ values were obtained fluorometrically and by an HPLC-based method when hexanal and all-*trans*-retinaldehyde (atRAL) were used as substrates, respectively. The reaction mixture was incubated in the corresponding reaction buffer<sup>71</sup> for 20 min at 37°C, before adding NAD<sup>+</sup> as cofactor, NADH as internal standard and the corresponding substrate. $k_{inact}/K_i$ values and the affinity labelling type were obtained by $k_{obs}$ vs [I] plot. Experimental values are the mean ± standard error.

To assess label specificity, we plotted *kobs versus* [I] and calculated the *kinact/Ki* value, representing the apparent second-order bimolecular rate constant (Table 1 and Supplementary Table S1). ABD0171 displayed the highest *kinact/Ki* values against ALDH1A3, showing specific affinity-labeling for this isoform, as indicated by the hyperbolic *kobs versus* [I] plot (Fig. 4a). In contrast, DIMATE showed specific affinity-labelling for ALDH1A1 (Fig 4a) but non-specific labeling for ALDH1A2 and ALDH2 (Fig. 2a and Supplementary Fig. 2a). At low concentrations (0-50 µM), DIMATE interacted specifically with ALDH1A3 (Supplementary Fig. 2b), while at higher concentrations, it acted as a non-specific affinity-label (Fig. 4a).

We confirmed the irreversible binding of DIMATE and ABD0171 to ALDH1A3 using electrospray ionization time-of-flight mass spectrometry (ESI-TOF-MS). Under varying voltage and temperature conditions, treatment with 167 µM DIMATE (1:10, ALDH1A3: inhibitor ratio) induced a complete shift in ALDH1A3 subunit mass resulting in species approximately 185 Da larger than the native protein subunit (Fig. 4b). We observed similar results with ABD0171, which generated species with a molecular mass increase of 333 Da compared to the untreated sample. At 50 µM (1:3, ALDH1A3: inhibitor ratio), DIMATE specifically modified ALDH1A3, leading to partial alteration of the native protein (Supplementary Fig. 2c). The mass spectra suggest irreversible modification with one DIMATE or ABD0171 molecule per protein subunit, corroborating our inhibition kinetics findings.

### ABD0171 induces death receptor-mediated extrinsic apoptosis and sensitizes basal subtype cells to anoikis by inhibiting key signaling pathways

Previous reports showed that DIMATE was able to induce cell death in p53-mutated cells^34,35^. To investigate thoroughly the mechanism of cell death of ABD0171 and DIMATE in TNBC cell lines, we used the MDAMB231 (Basal B, p53-mutated), MDAMB468 and HCC70 cells (Basal A, defective and wild-type p53, respectively). Annexin V/PI staining indicated that both compounds induce apoptosis at equimolar concentrations (Fig. 5a). Apoptosis triggered by ABD0171 and DIMATE was characterized by reduced expression of FLICE-inhibitory protein (c-FLIP), decreased of cellular X-linked inhibitor of apoptosis proteins (cIAP2 and XIAP), and activation of initiator caspase-8. This sequence of events is indicative of the extrinsic apoptosis pathway activation. Consequently, activated forms of effectors caspase-7, caspase-3 and its cleavage product cPARP were prominently detected in all cell lines treated with ABD0171, and to a lower extent in cells treated with DIMATE (Fig. 5b). Activation of death receptors was accompanied by reactive oxygen species (ROS) generation (Supplementary Fig. 3a) and sustained activation of the oxidative stress-sensing JNK pathway and its downstream effector c-Jun, for both compounds (Fig. 5b). Stress activation of p38 was transient, with DIMATE-treated cells showing greater prominence. Both extrinsic apoptosis and JNK pathway activation can influence inflammation and the metastatic potential of TNBC through several mechanisms. Therefore, we further investigate the effects of ABD0171 and DIMATE on cell migration, invasion and anoikis, which are critical processes in the metastatic cascade. At sub-lethal concentrations, ABD0171 and DIMATE attenuated migration and significantly reduced the invasion capacity of MDAMB231, MDAMB468, and HCC70 cells (Fig. 5c and Supplementary Fig. 3b). Similarly, the compounds significantly diminished the ability of these cell lines to resist anoikis in a dose-dependent manner after 24 hours (Fig. 5d and Supplementary Fig. 3b). Intriguingly, Basal A MDAMB468 and HCC70 cell lines displayed a shift from ALDH1A3 to ALDH1A1 under non-anchored conditions (Fig. 5e), becoming more evident by day 10, when cells formed spheroids with distinct morphological organization. This shift may reflect the acquisition of EMT-like features, as ALDH1A1 has been previously associated with mesenchymal traits and stemness, which are often enriched in spheroids formed by TNBC cell lines^36^.

**Figure 5.**
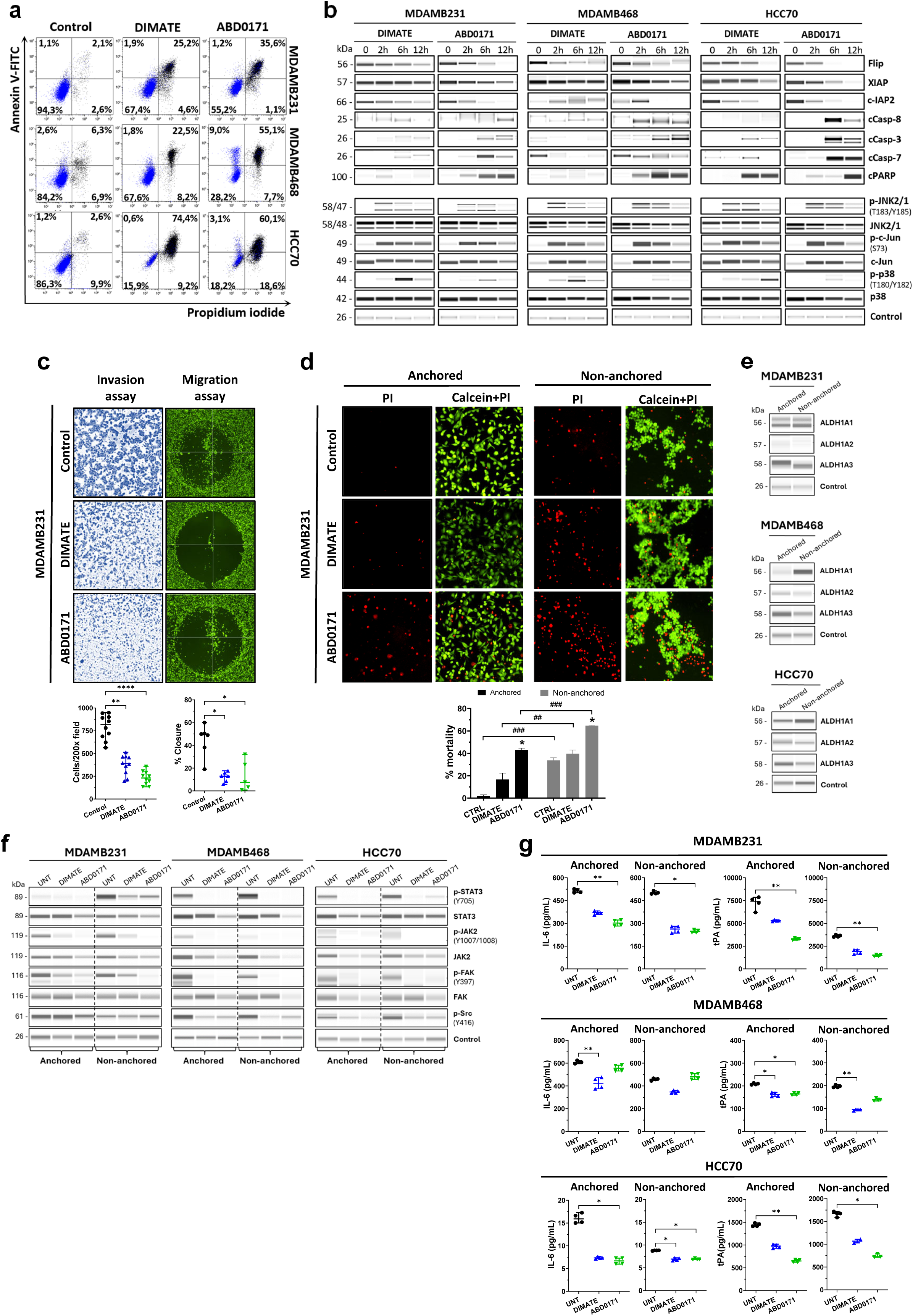
Effects of ALDH inhibitors ABD0171 and DIMATE on cell viability, invasion, migration and anoikis in triple-negative breast cancer (TNBC) cell lines. **(a)** Flow cytometry analysis of early and late apoptosis in TNBC cells treated with ABD0171 or DIMATE for 12h, using Annexin V/PI staining. The lower-left quadrant indicates healthy cells, while the lower-right and upper-right quadrants show early and late apoptotic cells, respectively. **(b)** Protein analysis using the WES (Western enhanced sensitivity) capillary electrophoresis system, illustrating the activation of the extrinsic apoptotic pathway and the JNK (c-Jun N-terminal kinase)/ cJun signaling pathway in response to equimolar concentrations of ABD0171 and DIMATE. **(c)** Representative images from invasion and migration assays in MDAMB231 cells treated with ABD0171 or DIMATE for 24-48h. Scatter plots (bottom) display quantification from two independent experiments (median ± standard deviation). Statistical analysis was performed using Kruskal-Wallis test. **(d)** Evaluation of anoikis resistance in MDAMB231 cells cultured on anchored and non-anchored (poly-HEMA-coated) surfaces and treated with either vehicle control, DIMATE or ABD0171 for 24h. Representative images of cells stained with calcein-AM and propidium iodide are shown, illustrating cell viability and death. The resazurin-based toxicity assay was used to quantify cell death, with results presented as median values ± standard deviation from triplicate experiments (bottom). Statistical significance was determined using an unpaired t-test with Welch’s correction. **(e)** WES capillary immunoelectrophoresis analysis of ALDH1A1, ALDH1A2 and ALDH1A3 protein expression levels in cells subjected non-anchored culture conditions for 10 days**. (f)** WES capillary immunoelectrophoresis analysis showing the activation STAT3 and associated proteins involved in its signalling pathway following 24h of treatment with ABD0171 or DIMATE. **(g)** Quantification of IL-6 and tPA protein levels by ELISA in TNBC cell lines, either untreated and treated with DIMATE and ABD0171 for 14h, under both anchored and non-anchored conditions.

As early as 24 hours, we observed robust activation of STAT3 in all cell lines under non-anchored conditions (Fig. 5f). The observation that ABD0171 and DIMATE reduce anoikis resistance in these cells suggested that both inhibitors could disrupt oncogenic reprogramming mediated by STAT3, triggered by detachment from the extracellular cell matrix. Consistently, ABD0171 and DIMATE simultaneously suppressed tPA, Src/FAK and IL6/JAK/STAT3 signaling pathways in anchored and non-anchored conditions, with ABD0171 demonstrating a markedly stronger effect (Fig. 5f, g). In contrast, DIMATE elicited pro-survival activation of AKT in MDAMB468 cells and ERK in HCC70 cells, suggesting potential compensatory mechanisms that were not observed with ABD0171 (Supplementary Fig. 3c).

### ABD0171 reduces TNBC cell proliferation and metastasis in mouse models, with demonstrated *in vivo* safety

Before evaluating *in vivo* efficacy, we assessed potential systemic, hepatic, and/or renal toxicity following repeated administration of ABD0171 and DIMATE in healthy animals. We administered equimolar doses of both compounds (30 and 40 mg/kg for ABD0171; 15 and 20 mg/kg for DIMATE) three times per week over 29 days. Neither 30 nor 40mg/kg of ABD0171, nor 15 mg/kg of DIMATE caused body weight loss in healthy animals (Supplementary Fig. 4a and Supplementary Data 3E) or produced visible signs of adverse events. In these groups we did not observe any significant alterations in serum biochemical parameters, including liver enzymes (alkaline phosphatase, ALP; alanine aminotransferase, ALT; gamma-glutamyl transferase, GGT), bile acids (BA), total bilirubin (TBIL), albumin (ALB), renal markers (blood urea nitrogen, BUN; creatinine, CRE), or electrolytes (Na⁺, K⁺, Cl⁻). Other factors such as cholesterol (CHOL), glucose (GLU), calcium (Ca²⁺), phosphate (PHOS), and total CO₂ (tCO₂) also remained unaffected, suggesting no impairment of hepatic or renal function. Hematological analysis showed no significant changes in red blood cell count (RBC), hemoglobin (Hb) levels, leukocyte numbers, or other key hematological parameters (Supplementary Data 3F). However, in the group treated with 20 mg/kg of DIMATE, we observed a transient decrease in body weight (Supplementary Fig. 4a and Supplementary Data 3E), accompanied by a moderate increase in tCO₂ levels (p=0.003) compared to controls, remaining within a non-critical range (Supplementary Data 3F).

We next evaluated the anticancer efficacy of ABD0171 and DIMATE *in vivo*, using two different TNBC orthotopic mouse models: the MDAMB231-GFP xenograft and the immunocompetent 4T1-Luc_M3 model. In both models, treatment with ABD0171 was well tolerated without observed weight loss (Supplementary Fig. 4b, c). Contrarily, DIMATE administration caused noticeable loss of weight in immunocompetent mice and lethal acute toxicity in 6 animals in the immunosuppressed MDAMB231-GFP model towards the end of the study (Fig. 6a, Supplementary Fig. 4b, c and Supplementary Data 3G). Both compounds resulted in significant inhibition of tumor growth (Fig. 6a, b). On day 30 of the experiment, treated mice bearing MDAMB231-GFP tumors showed tumor growth reductions of 54% and 32% with 40 mg/kg ABD0171 and 20 mg/kg DIMATE, respectively, compared to controls (p < 0.0001, p = 01710) (Fig. 6a). Similarly, both compounds reduced 4T1-Luc_M3 tumor growth by 14% and 20% on day 24 of the experiment (p = 0.0430, p = 0.0004, respectively) (Fig. 6b and Supplementary Data 3H).

**Figure 6.**
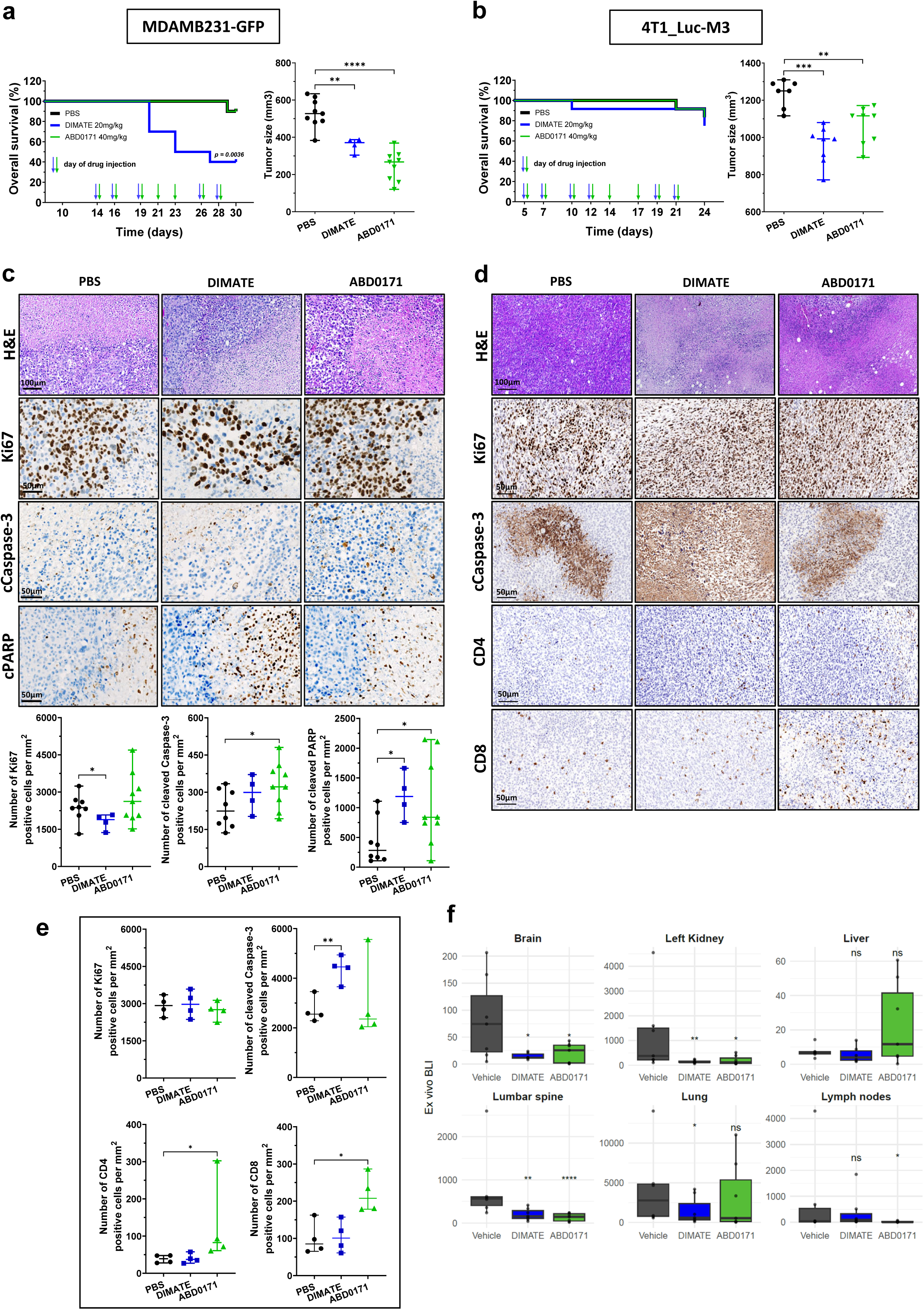
ALDH inhibitors reduce tumor growth in immunocompromised and immunocompetent TNBC cancer models. **(a, b)** Assessment of survival and antitumor efficacy of DIMATE and ABD0171 in two murine TNBC models. SCID-CB17 mice bearing MDAMB231-GFP tumors and BALB/c mice with 4T1-Luc_M3 tumors were treated intraperitoneally with equimolar concentrations of DIMATE (20 mg/kg) or ABD0171 (40 mg/kg), administered three times per week for 3 weeks. Control groups were treated with compound’s vehicle. Tumor growth inhibition is shown in the right panel, and overall survival rates are presented in the left panel. **(c, d, e)** Representative immunohistochemistry images for the indicated antibodies are provided. Quantification of positive cells was performed using QuPath. **(f)** Quantification of metastatic spread in the indicated organs, determined by luminescence intensity through bioluminescence imaging.

Histopathological analysis of MDAMB231 tumor tissues revealed neoplastic proliferation with pleomorphic cells, absence of glandular or ductal structures, poor vascularization, and extensive necrosis (Fig. 6c). Treatment with DIMATE and ABD0171 resulted in a significant increase in cleaved caspase-3 and cleaved PARP-positive cells compared to controls, indicating enhanced apoptosis (Fig. 6c and Supplementary Data 3H).

Similarly, 4T1-M3 tumor tissues displayed disorganized architecture, dense cellularity, pleomorphic cells with scant cytoplasm, and high Ki67 positivity, consistent with elevated proliferation (Fig. 6d). While DIMATE-treated tumors showed increased cleaved caspase-3 positivity, this effect was not observed with ABD0171 treatment. However, ABD0171-treated tumors demonstrated heightened T-cell infiltration, predominantly CD8+ and, to a lesser extent, CD4+ T cells, suggesting a potential immunomodulatory effect (Fig. 6d, e and Supplementary Data 3H). Complementing these primary tumor effects, treatment with ABD0171 and DIMATE significantly reduced metastasis to lymph nodes, brain, lumbar spine, and kidneys in this model (Fig. 6f and Supplementary Data 3I), supporting the antimetastatic activity observed in the functional *in vitro* studies and highlighting the therapeutic potential of these compounds in targeting both primary tumor growth and metastatic spread.

## DISCUSSION

Drug resistance and the persistence of breast cancer cells with stem/progenitor properties remain significant challenges in treating aggressive breast cancer^37,38^. ALDH1 activity has been closely associated with stemness features and plays crucial roles in driving this therapeutic resistance^10,11,37–39^. In this study, we thoroughly characterized the expression diversity of the ALDH1A subfamily across molecular subtypes and cellular populations in breast cancer, uncovering distinct profiles of *ALDH1A1*, *ALDH1A2*, and *ALDH1A3* in both patient-derived tumors and breast cancer cell lines. Our analysis highlights the varied roles that these isoforms may play within different pathological entities and the tumor microenvironment, emphasizing their potential as therapeutic targets.

One of the key findings of this study is the predominant expression of *ALDH1A3* in the Basal-like molecular subtype of breast cancer, where it clusters with Basal-like gene markers in the PAM50 signature^22,23^. The elevated expression of *ALDH1A3*, primarily attributed here to the epithelial cancer cell compartment within tumors, is consistent with the significantly higher levels of ALDH1A3 observed in the Basal A subtype of TNBC cell lines, which are also characterized by epithelial-like features^26^. These results complement and expand prior studies linking ALDH1A3 expression to aggressive TNBC phenotypes and poor clinical outcomes^15,17,18,40^. TNBC and Basal-like breast cancers often exhibit overlapping morphological characteristics and biological aggressiveness; however, they are not entirely identical entities^28,41–43^. Basal-like TNBC tumors exhibit distinct molecular alterations (e.g. TP53 mutations and increased mitotic activity), unique patterns of organ-specific metastases as well as worse survival outcomes compared to TNBC tumors that are not Basal-like^42,43^. Additionally, other breast cancer subtypes, including specific HER2-enriched tumors, certain luminal B tumors, and the majority of claudin-low phenotype tumors, can exhibit molecular and histological characteristics similar to Basal-like cancers^24,28^.

Interestingly, recent findings from a comprehensive multi-dimensional atlas of breast tissues identified a distinct subset of cells termed basal luminal alveolar cells^44^. These epithelial cells accumulate with age, exhibit low lineage fidelity, and express high levels of stem/progenitor markers ALDH1A3 and CD73. Notably, their gene expression signatures strongly associate with Basal-like breast cancers, aligning with our observations and suggesting that they may represent a critical ALDH1A3-enriched population implicated in tumor initiation. These findings highlight the potential of ALDH1A3 as both a Basal-like biomarker surrogate and a therapeutic target. While further investigation is necessary, incorporating ALDH1A3 into immunohistochemistry strategies could enhance the sensitivity of existing biomarker panels for Basal-like TNBC, which currently rely on basal marker staining (cytokeratin 5/6, -14, or -17, and/or EGFR) in combination with hormone receptor and HER2 negativity, achieving a reported sensitivity of 76%^45^.

Our single-cell analysis also provides valuable insights into the expression of *ALDH1A1*, a marker associated with breast cancer stem cells and poor prognosis^12^, and *ALDH1A2* isoforms within breast tumors. Unlike *ALDH1A3*, *ALDH1A1* and *ALDH1A2* were less prominently expressed in cancer epithelial cells, displaying distinct cellular distribution profiles. ALDH1A1 was predominantly enriched in myeloid cells, while ALDH1A2 was primarily expressed in endothelial cells. Although the three isoforms were detected in myCAF-like cells, only ALDH1A1 was significantly expressed in this relevant subtype of cancer-associated fibroblasts known for its contribution to mechanical support, extracellular matrix remodeling, and stromal stiffness, facilitating tumor invasion. An intriguing aspect of our findings is the high expression of *ALDH1A1* and *ALDH1A2* in claudin-low breast cancers and in cell lines resembling these tumors. This observation aligns with a recent study that identified *ALDH1A1* enrichment in a specific claudin-low subgroup of breast tumors, characterized by high stemness scores, immune cell infiltration, and strong epithelial-to-mesenchymal transition (EMT) marker expression^2^. The authors suggested that the observed *ALDH1A1* expression could partly result from infiltrating immune cells. Furthermore, our *in vitro* experiments demonstrated that breast cancer cells can transition from ALDH1A3 to ALDH1A1 protein expression under EMT-inducing conditions, such as anchorage-independent growth, highlighting a plasticity in ALDH1 isoforms expression in response to tumor microenvironment cues. The prominent expression of ALDH1A1 and ALDH1A2 in different stromal populations and in claudin-low tumors, known for their stromal richness, suggests distinct functional roles within these tumors. ALDH1A1 likely contributes to myeloid-associated tumor immunity by modulating the metabolic adaptation of myeloid cells within the tumor microenvironment^30,46^ and promoting the mesenchymal traits characteristic of these primitive tumors. These findings highlight the need for further investigation into the specific roles of ALDH1 isoforms in aggressive breast cancer phenotypes and emphasize the importance of further exploring the role of ALDH1A1 in immunosuppression, ALDH1A2 in angiogenesis, and the broader contribution of ALDH1 isoforms on matrix remodeling within the tumor microenvironment.

The profiling of ALDH1A isoforms and the therapeutic hypotheses that they prompt guided our efforts to develop novel ALDH1A inhibitors, designed as structural analogues of DIMATE, a clinical-stage ALDH inhibitor^47^. Among these, ABD0171 emerged as a potent irreversible inhibitor with specific affinity labeling properties for ALDH1A3. Belonging to the series V of analogs, ABD0171 demonstrated enhanced efficacy against ALDH1A3-expressing Basal-like cancer cells compared to DIMATE. Structure-activity relationship analyses revealed the critical role of aryl substitutions in series V analogs for enhancing inhibitory activity, enabling ABD0171 to achieve GI50 values below 10 µM in most Basal A and Basal B cell lines.

A key challenge in translating novel ALDH1A3 inhibitors into the clinical setting lies in achieving selectivity while sparing ALDH2, the closest related enzyme within the ALDH1A subfamily based on evolutionary divergence^48^. Unlike the cytoplasmic ALDH1A isoforms, ALDH2 is localized to mitochondria and plays an essential role in hematopoiesis, the prevention of mutagenesis in somatic tissues and cardioprotection through its enzymatic activity, which mitigates oxidative stress and clears toxic endogenous acetaldehyde and formaldehyde^49^. ABD0171 exerts only marginal inhibitory effects on ALDH2 and demonstrates low potency against ALDH3A1, an isoform that constitutes 10–40% of the water-soluble protein in the mammalian cornea and plays a crucial role in protecting this tissue from oxidative stress^50^. This selective inhibition of ALDH1A3, combined with favorable cytotoxicity profiles, underscores the potential of ABD0171 to overcome the safety challenges observed with broader ALDH inhibitors, making it a highly promising therapeutic candidate for further clinical development.

In comparative functional studies, ABD0171 exhibited higher predictive cytotoxicity in TNBC models, as determined through linear mixed modeling. It also demonstrated a stronger induction of death receptor-mediated apoptosis in TNBC cell lines compared to DIMATE. These apoptotic events were associated with elevated ROS levels and sustained activation of the oxidative stress-sensing JNK pathway (JNK/cJun). The differences in apoptotic effector activation between ABD0171 and DIMATE may be attributed to the concurrent activation of prosurvival pathways, such as AKT and ERK, and the transient activation of p38 observed prominently in DIMATE-treated cells. While AKT and ERK signaling are well known to promote cell survival and counteract apoptosis, the transient activation of p38 may further induce a prosurvival stress response that mitigates caspase activation. This mechanism has been previously reported under chemotherapy-induced stress conditions, such as those induced by Paclitaxel and doxorubicin in TNBC cells^51,52^. This further underscores the differential capacities of ABD0171 and DIMATE to modulate stress-response pathways in TNBC models.

*In vivo*, both ABD0171 and DIMATE significantly inhibited tumor growth in MDA-MB-231 xenografts and syngeneic 4T1-Luc_M3 TNBC models. ABD0171 demonstrated superior tolerability, reflected in its favorable safety profile, with minimal systemic, hepatic, or renal toxicity. Notably, ABD0171 uniquely enhanced cytotoxic CD8+ T cell infiltration into the tumor microenvironment, an observation that warrants further investigation, particularly in TNBC, where the tumor microenvironment is predominantly immunosuppressive, often compromising responses to immunotherapies. This finding raises the possibility of ABD0171 not only as a direct antitumor agent but also as a potential modulator of tumor immunity, offering potential benefits in combination with immune checkpoint inhibitors. Moreover, ABD0171 was more effective than DIMATE in reducing tumor burden and metastasis to distant organs, including lymph nodes and the lumbar spine, while both compounds exhibited comparable antimetastatic activity in the brain and kidney.

These findings align with our *in vitro* studies, where sub-lethal concentrations of ABD0171 and DIMATE suppressed key metastatic traits such as migration, invasion, and anoikis resistance in TNBC cells by disrupting the tPA, Src/FAK, and IL6/JAK/STAT3 signaling pathways. The modulation of these pathways through ALDH1A3 inhibition underscores its pivotal role in TNBC progression. tPA, a key mediator of aggressive breast cancer phenotypes, has been shown to promote invasion and lymph node metastasis through its regulation by ALDH1A3-derived RA, the principal product of ALDH1A enzymatic activity and a major regulator of gene expression in tumor biology^16^. RA is reported also to modulate the expression of Src activators such as integrins and EGFR, which are frequently upregulated in TNBC^53^. These activators, together with elevated tPA levels, converge to activate Src kinase and FAK, driving cytoskeletal remodeling and cell motility, hallmarks of metastatic dissemination54,55.

RA is also reported to influence cytokine production, notably IL6, a critical activator of the JAK/STAT3 pathway56. This central axis governs multiple oncogenic processes, including epithelial-to-mesenchymal transition (EMT), tumor proliferation, chemoresistance, immune evasion, and metastasis57,58. Aberrant activation of STAT3, observed in approximately 50–70% of TNBC cases, amplifies a plethora of oncogenic signaling, making it a key driver of TNBC pathogenesis and a significant therapeutic target in breast cancer research59–61. Our findings underscore the potential of ALDH1A3 inhibition to disrupt a broad spectrum of signaling networks integral to Basal-like or TNBC subtypes in general and suggest additional therapeutic potential in combination strategies to overcome immune evasion in aggressive breast cancers.

In conclusion, the distinct expression patterns of ALDH1A3 across breast cancer subtypes and tumor cell populations points to its association with a specific subset of epithelial cells, characterized by unique properties that strongly contribute to disease initiation and progression. Furthermore, the predominance of ALDH1A1 and ALDH1A2 in specific stromal cells provides valuable insights into the complex biology of breast cancer, highlighting the need to address both tumor genetic heterogeneity and the intricate tumor-stroma interactions when designing ALDH-targeted therapies.

The development of selective ALDH1A3 inhibitors, such as ABD0171, represents a promising approach to tackle the therapeutic challenges linked to aggressive breast cancer subtypes, including TNBC and Basal-like cancers. By targeting key signaling pathways, ABD0171 has the potential to significantly enhance patient outcomes in these subtypes, where effective treatment options remain limited. Future research should focus on evaluating combinatorial therapies and deepening our understanding of the roles of ALDH1A in tumor immunity and resistance mechanisms. These steps will be critical to unlock the full clinical potential of ALDH1A inhibition as a transformative strategy in breast cancer therapy.

## METHODS

### Public data-based transcriptomic profiling of breast cancer cell lines and tumors

#### Breast cancer cell lines

CCLE 2019 bulk RNA-seq data (raw read counts) for 50 human breast cancer cell lines were retrieved from the Dependency Map portal (DepMap)^25^, where reads had previously been aligned to the reference genome (GRCh37.p13, GENCODE release 19). Genes with low expression were filtered out, retaining only those with sufficiently high counts for further statistical analysis. Using the edgeR package^62^, normalization factors were calculated to scale the raw library sizes via the trimmed mean of M-values (TMM) method. Log_2_ counts per million (CPM) values were then computed as a measure of gene expression levels. The molecular subtypes of the cell lines were kept as annotated in DepMap, ensuring consistency with existing classification standards. For unsupervised clustering, the log_2_ expression of 59 genes, which included the well-known PAM50 signature^22,23^, *CHD1, CLDN3, CLDN4, CLDN5, CLDN7, OCLN*, and *ALDH1A* isoforms (*ALDH1A1, ALDH1A2, ALDH1A3*), was used as input. This analysis was conducted using the partitioning around medoids (PAM) algorithm set to k=4 (Supplementary Data 1B and Fig. 1c). Subsequently, the median expression of these 59 genes was calculated for each of the four groups, based on the standard molecular classification of breast cancer cell lines: Basal A, Basal B, HER2-enriched, and Luminal.

#### Breast tumor patient samples

mRNA (bulk) gene expression data from three breast cancer studies were retrieved from cBioPortal (https://www.cbioportal.org): Breast Invasive Carcinoma (TCGA, Cell 2015)^21^, Breast Invasive Carcinoma (TCGA, PanCancer Atlas)^20^, and Breast Invasive Carcinoma (TCGA, Firehose Legacy). After merging the individual patient samples, a total of 1103 unique samples were selected across the three studies. As described above, using the edgeR package, normalization factors were calculated to scale the raw library sizes via the trimmed mean of M-values (TMM) method. Log_2_ counts per million (CPM) values were then computed as a measure of gene expression levels. Next, the log_2_ expression of 59 genes was used as input for unsupervised clustering using the PAM algorithm with k=5 (Supplementary Data 1A and Fig. 1b). Following this, the median expression of these 59 genes was calculated for each of the five groups, based on the molecular classification: Basal-like, Claudin-low, HER2-enriched, Luminal A, and Luminal B.

#### Single-cell of breast cancer cell lines

Pre-processed scRNA-seq data from 32 cell lines, initially comprising 35,276 cells, was retrieved from: https://figshare.com/articles/dataset/Single_Cell_Breast_Cancer_cell-line_Atlas/15022698^29^. Subsequently, genes with high relative expression per cell were filtered out, and the analysis focused on a subset consisting only of cell lines classified as Basal A or Basal B, which represents 16 cell lines encompassing a total of 20,485 cells after filtering. Further data processing including normalization, dimensionality reduction, and clustering was conducted with the Seurat R package (v4.4)^63^. Normalization was performed using the sctransform v2 method with the glmGamPoi package^64^. The FeaturePlot and VlnPlot functions from Seurat were used to visualize specific gene expressions across identified clusters.

#### Single-cell of breast tumor patients

Pre-processed scRNA-seq data from breast cancer patients, comprising 26 individual patient samples and a total of 100,064 cells, was retrieved from https://singlecell.broadinstitute.org/single_cell/study/SCP1039^30^. Further data processing including normalization, dimensionality reduction, and clustering was conducted with the Seurat R package (v4.4)^63^. Prior to data integration, individual patient samples were grouped according to their molecular classification: Basal-like, HER2-enriched, Luminal B/HER2+, Luminal A, and Luminal B. Normalization was performed using the sctransform v2 method with the glmGamPoi package. Additionally, to remove potential confounding sources of variation, the mitochondrial mapping percentage was regressed during normalization. Data integration was performed using the SCT method and 3000 features for anchoring. The number of principal components was set to 30. The FeaturePlot function from Seurat was used to visualize specific gene expressions across identified clusters. Visualization was conducted using ggplot2^65^, GGally^66^, complexHeatmap^67^ and Seurat^63^ R packages.

### Chemical synthesis of compounds

Various structural derivatives of DIMATE were synthesized according to the following scheme:

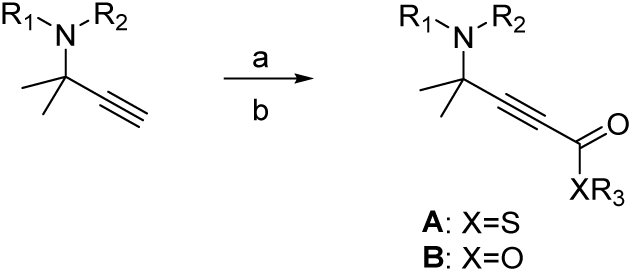

Aminothiolesters synthesis: **a:** nBuLi, THF, -70°C to 0°C – b: COS, -70°C to 0°C / R3I, 0°C [method 1] or CO2, 0°C to 20°C / isobutyl chloroformate, 0°C to 20°C/ NaSMe, 20°C [method 2]

The process and experimental details for the preparation of compounds **A** or **B** have been described previously^32,33,68,69^. Briefly, aminothiolesters **A** were obtained by deprotonation of corresponding acetylenic amines with n-Butyllithium. The resulting acetylides were treated with carbonyl sulfide (COS) then with alkyl iodide [method 1]. An alternative process involved the reaction of lithium acetylide intermediate with carbon dioxide followed by treatment with isobutyl chloroformate and finally with sodium methanethiolate (NaSMe) [Method 2]. CEY1410 (R1, R2=Me, X=O, R3=H) was performed by treatment of lithium acetylide intermediate by carbon dioxide followed by an acidic work-up. Corresponding alkylesters **B** (R3=Me, Et) were obtained by Fisher esterification of CEY1410.

### Protein expression and purification

*Escherichia coli* BL21 (DE3) pLysS strain was transformed with the plasmid pET-30 Xa/LIC that encoded ALDH1A1, ALDH1A2, ALDH1A3, ALDH3A1 or ALDH2 fused to the N-terminal His_(6)_ tag. The ALDH isoforms were expressed and purified as previously described^70^ using a nickel-charged chelating Sepharose™ Fast Flow 5 mL column (HisTrap) and an ÄKTA™ FPLC purification system.

### Enzymatic activity assay

Standard activities were measured prior to each inhibition experiment by using hexanal (for ALDH1A and ALDH2) or 4-nitrobenzaldehyde (for ALDH3A1) as a substrate as described^70^. DTT was removed from the reaction buffer and the pH was adjusted at 7.2. Thus, 50 mM HEPES, 0.5 mM EDTA, pH 7.2 was used for ALDH1A1, ALDH1A2 and ALDH2; 50 mM HEPES, 30 mM MgCl_2_, pH 7.2 was used for ALDH1A3; and 50 mM Tris/HCl, pH 7.2 was used for ALDH3A1. Enzymatic activity assays were performed at 37°C using a Cary Eclipse fluorescence spectrometer.

### Inhibitor screening for human ALDH1A, ALDH3A1 and ALDH2 isoforms and determination of the IC_50_ values

Single-point analyses of enzymatic activity at 10 µM of inhibitor were performed by the fluorimeter methodology described previously^70^. Enzymatic inhibition assays were performed at 37°C, in a final volume of 1 mL and in an appropriate reaction buffer. All compounds tested as inhibitors (including DIMATE and ABD0171, Fig. 3b) were dissolved in ethanol and assayed in a final concentration of 1% ethanol (v/v) using hexanal (for ALDH1A and ALDH2) and 4-nitrobenzaldehyde (for ALDH3A1) as a substrate. The inhibitor effect was checked at both saturating and *K_m_* concentrations of substrate^71^. Usually, we performed the inhibition experiments at saturating concentrations of substrate for irreversible and non-competitive inhibitors but substrate concentrations at *K_m_* values were also used to identify possible competitive inhibition. For this reason, the inhibitory effect of compounds was tested at two substrate concentrations: 10 and 0.25 µM hexanal for ALDH1A1, at 250 and 10 µM hexanal for the remaining ALDH1A and ALDH2 enzymes and at 250 and 5 µM 4-nitrobenzaldehyde for ALDH3A1. The concentration of enzyme was kept from 50- to 100-fold lower than that of the substrate. The reaction mixture containing enzyme and inhibitor in the appropriate reaction buffer was incubated for 20 min at 37°C, before adding the corresponding NAD^+^/NADP^+^ cofactor, NADH/NADPH internal standard and substrate (hexanal/4-nitrobenzaldehyde). The percentage of remaining activity was calculated from the ratio of activity at a given inhibitor concentration to the control activity with 1% ethanol (v/v) without inhibitor. The IC_50_ values were calculated at saturating concentrations of substrate by fitting the initial rates to the appropriate equation using GraFit 5.0 (Erithacus Software) and the values were given as the mean ± standard error.

### Time-dependent inhibition assay and determination of *k_obs_* and *k_inact_/K_i_* parameters

Different concentrations of DIMATE and ABD0171 were incubated in a range of time including 0, 2, 5, 10, 15, 20, 30, 45, 60, 90, 120, 180 min at 37°C with ALDH enzymes (ALDH1A and ALDH2) in the corresponding reaction buffer, before to the addition of NAD^+^ cofactor, NADH internal standard and hexanal at saturating concentrations and at concentrations near *K_m_* values. The corresponding inhibitor solutions were prepared to be assayed in a final concentration of 1% ethanol (v/v). The concentration of enzyme was kept from 50- to 500-fold lower than that of the substrate for all enzymatic assays and 500 µM NAD^+^ was added after preincubation time. The inhibitory effect of compounds was tested using the fluorometric assay, at 10 and 0.25 µM hexanal for ALDH1A1, at 250 and 10 µM hexanal for ALDH1A2, ALDH1A3 and ALDH2 enzymes, to determine the effect of substrate concentrations on the inhibition. In all cases, *k_obs_* was obtained at saturating concentrations of substrate by fitting the data using GraFit 5.0 to the following equation: *v_t_ = v_i_ ꞏ exp (*−*k_obs_ ꞏ t)*, where *v_t_* is the measured steady state velocity and *v_i_* is the initial velocity at short preincubation times (*t)*. Finally, the dependence of *k_obs_* on inhibitor concentration was represented to obtain the *k_inact_*/*K_i_* rate constant, which is the most reported parameter to describe time-dependent inhibitors^72^.

### Non-competitive inhibition of DIMATE and ABD0171 against ALDH3A1 and determination of *K_i_ value*

The inhibitory effect of DIMATE and ABD0171 against ALDH3A1 was analyzed fluorometrically using 4-nitrobenzaldehyde as a substrate, 1 mM NADP^+^ as a cofactor and 5 µM NADPH as an internal standard in the corresponding reaction buffer. Activity assays to determine the type of inhibition and *K_i_* values were performed using various substrate concentrations at fixed inhibitor concentrations (0, 50, 250 and 500 µM DIMATE and 0, 100, 250 and 500 µM ABD0171). The data of enzymatic activities were fitted to the Michaelis-Menten equation using GraFit 5.0, to determine the values of *K_m_* and *V_max_*. Then, results were fitted to the equations of the different types of enzymatic inhibition and *K_i_* values were determined choosing the type of inhibition yielding the best fit. Values were expressed as the mean ± standard error.

### HPLC-based assay for the inhibition characterization using all-*trans*-retinaldehyde as a substrate

To evaluate the retinaldehyde dehydrogenase activity of human ALDH1A isoforms, an HPLC method was used in presence of all-*trans*-retinaldehyde as a substrate and DIMATE and ABD0171 as inhibitors. Stock solutions of retinoid substrate were stored in ethanol. Working solutions of retinoids were prepared by 10-min sonication in the presence of equimolar delipidated BSA^73^. The actual concentration of solubilized retinoid was determined based on the corresponding molar absorption coefficient (ε_370_ = 29,500 M^−1^ꞏcm^−1^). The enzymatic reaction was carried out for 15 min at 37°C in a final volume of 500 µL, after 20-min preincubation with different concentrations of DIMATE and ABD0171 and using a saturating concentration of cofactor (500 µM NAD^+^) in the corresponding reaction buffer. With the aim to measure the steady-state enzymatic activity, the concentration of enzymes was kept from 50- to 100-fold lower than that of the substrate for all enzymatic assays. The reaction was stopped with 1 mL cold hexane:dioxane:isopropanol (50:5:1, v/v/v), and retinoids were extracted by a two-step procedure with the same solvent mixture^74,75^. The aqueous phase was removed, and the organic phase was evaporated under a N_2_ stream. Retinoids were dissolved in 200 µL of hexane and analyzed by a modification of a published method^76^, as follows. Retinoids were separated by HPLC on a Novapak® Silica 4 µm, 3.9 x 150 mm in hexane/methyl-tert-butyl ether (96:4, v/v) mobile phase, at a flow rate of 2 mL/min, using a Waters Alliance 2695 HPLC. Elution was monitored at 370 nm and 350 nm for all-*trans*-retinaldehyde and all-*trans*-retinoic acid, respectively, using a Waters 2996 photodiode array detector. Quantification of retinoids was performed by interpolation of HPLC peak areas into a calibration curve of known retinoid concentration (data not shown). IC_50_ and *k_obs_*parameters were calculated by fitting the initial rates to the appropriate equation using GraFit 5.0 and values were given as the mean ± standard error. All compound manipulations were performed under dim red light to prevent retinoid photoisomerization.

### Electrospray Ionization Time-of-Flight Mass Spectrometry (ESI-TOF-MS) assays

ALDH1A3 wild type was stored at −20°C in 100 mM ammonium bicarbonate, pH 7.5, at 3.75 mg/mL (16.74 µM). The concentration of DIMATE and ABD0171 stock was prepared 100-fold higher in order to work at 1% EtOH (v/v). Fresh protein samples were incubated with the corresponding compound for 45 min at 37°C. The molecular weight of 16.7 µM ALDH1A3 formed after 45 min-incubation with 167 µM DIMATE, 50 µM DIMATE and 167 µM ABD0171 was determined by Electrospray Ionization Time-of-Flight Mass Spectrometry (ESI-TOF-MS) on a Micro TOF-Q instrument (Bruker Daltonics, Bremen, Germany), connected to a Series 1200 HPLC Agilent pump and controlled by the Compass Software. Samples were analyzed under neutral conditions (pH 7.0), using as running buffer a mixture of acetonitrile:ammonium acetate (15 mM) solution (20:80, v/v). For each analysis, 20 µL of protein solution were injected at 30-40 µLꞏmin^−1^ and analyzed under the following conditions: capillary counter-electrode voltage, 3500-5000 V; dry temperature, 90-110°C; dry gas, 6 Lꞏ min^−1^; m/z range 800-3000.

### Cell lines and culturing

The human breast cancer cell lines (Supplementary Data 3A) used in this study included DU4475, HCC1143, HCC1937, MDAMB231, MDAMB468, MDAMB453, SKBR3, MCF7, and BT474, obtained from the German Collection of Microorganisms and Cell Cultures GmbH (DSMZ), as well as HCC70, BT20, and MDAMB436, purchased from the American Type Culture Collection (ATCC). Cell lines HCC1419, HCC1569, HCC1954, MDAMB361, and T47D were generously provided by Dr. Ana Ruiz-Saenz (Dept. of Cell Biology, Erasmus Medical Center, Rotterdam, Netherlands), and Hs578T was kindly provided by Dr. Aida Zulueta-Morales (Dept. of Health Sciences, University of Milan, Italy).

The authenticity of cell lines was verified by polymorphic short tandem repeat (STR) profiling (Microsynth, Switzerland). All cell lines were cultured following the supplier’s recommended media and supplements and were maintained at 37°C in a humidified incubator with 5% CO₂.

### Cell cytotoxicity assay

Cells (0.5 x10^4^ cells/well) were seeded into 96-well cell culture plates and incubated with various concentrations of drugs (1–50μM) or control medium for 48 hours. The growth-inhibitory effect of the drugs was analyzed using an *in vitro*, resazurin-based toxicity assay, following the manufacturer’s instructions (Sigma-Aldrich, Darmstadt). The drug response was quantified by the half-maximal inhibitory growth concentration (GI_50_) and determined by non-linear regression analysis of log-dose/response curves. The cut-off value to define cell resistance was determined statistically (above GI50 geometric mean+ S.D.).

### 3D clonogenic assay

Patient-derived tumor xenografts (PDX) cells were plated in ultra-low attachment 96-well plates containing soft-agar medium and incubated with various concentrations of two drugs, ABD0171 and DIMATE (1-30µM) or control medium for 8 to 13 days, at 37°C and humidified air with 7.5% CO2. Colonies were stained with 2-(4-iodophenyl)-3-(4-nitrophenyl)-5-phenyltetrazolium chloride and counted with an automatic image analysis system (CellInsight NXT, Thermo Scientific or Bioreader 5000 V, BIO-SYS).

Data was collected from 11 models (7 TNBC, and 4 HER2-enriched), each treated with 9 different drug concentrations, along with control samples (untreated). Control values were calculated using the median across replicates and were used to normalize the response for each drug concentration. After normalization, the mean was calculated for two or three replicates per concentration, depending on the PDX model, to generate the final response values for each drug concentration. Dose-response curves were fitted for each model and drug using the dr4pl R package^77^, and the GI_50_ values were extracted from the fitted models.

### Linear mixed-effects model analysis

GI_50_ values, representing drug efficacy in both cell lines and PDX models, were selected for analysis. In the case of cell lines, molecular subtypes “Basal A” and “Basal B” were reclassified as triple-negative breast cancer (TNBC) to match the classification used for the PDX models, where only HER2-enriched and TNBC subtypes were available. A combined dataset of cell lines and PDX models was used to fit a linear mixed-effects model (LMM) using the lmerTest R package^78^. This model assessed the effects of drug (DIMATE or ABD0171), molecular subtype (TNBC or HER2-enriched), and their interaction on the observed GI_50_ values. The model included a random intercept to account for variability between individual models (both cell lines and PDXs). Satterthwaite’s approximation, provided by the lmerTest package, was applied to calculate degrees of freedom and p-values for the fixed effects. Both observed and predicted GI_50_ values were extracted for further analysis and interpretation.

### Apoptosis assays

Apoptotic cells were quantified using eBioscience Annexin V Apoptosis Detection Kit FITC (Thermo Scientific, Strasbourg, FR). Cells were incubated with Annexin V-FITC and propidium iodide (PI) according to the manufacturer’s instructions. Staining was performed for 15 minutes at room temperature, followed by analysis via flow cytometry. Annexin V-positive/PI-negative cells were classified as early apoptotic, while Annexin V-positive/PI-positive cells were considered late apoptotic or necrotic.

### Capillary electrophoresis immunoassay

For protein expression analysis, cells were lysed in RIPA buffer containing phosphatase and protease inhibitors (Calbiochem, San Diego, CA, USA). Total protein extracts were quantified and separated via capillary electrophoresis, using the WES Simple Western system and following manufacture’s specifications (Bio-techne, Minneapolis, USA). Immunodetection was performed using WES standard protocols^79^. Quantification of chemiluminescence signals and imaging were performed using Compass software for WES Simple western instruments (Bio-techne). Antibodies used in these studies are listed in Supplementary Table S2.

### Cell migration and invasion assays

Migration assays were performed using the Oris Cell Migration Assembly Kit-FLEX (PlatypusTechnologies, Madison, USA). Cells (5 × 10⁴ per well) were seeded into a 96-well plate containing an insert that created a defined, cell-free gap. After 24h of incubation to reach 100% confluence, ALDH inhibitors were added to the culture media, and the inserts were removed to allow migration. The cell-free area was monitored over 24h and quantified using the SpectraMax I3-Minimax station (Molecular Devices, USA) and ImageJ software^80^. Calcein AM staining (Fisher Scientific) was used for imaging at 0 and 24h. Data were obtained in triplicate for each condition.

For invasion assays^81^, cells (5–10 × 10⁴) were seeded onto ECM-coated filter membranes within transwell inserts (Corning, Lyon, FR). The lower chamber of a 24-well plate was filled with 600 μL of complete media, with or without ALDH inhibitors. After 24– 48h of incubation, non-migrated cells on the upper membrane surface were removed using cotton swabs. Migrated cells on the underside of the membrane were fixed in 70% ethanol, stained with 0.2% crystal violet, and quantified by manual counting under a microscope.

### Anoikis assay

Cells were cultured in 96-well plates coated with poly-HEMA (non-anchored conditions) or uncoated plates (anchored conditions) and treated with DIMATE or ABD0171 for 24h. Calcein AM and propidium iodide (PI) were sequentially added to each well, incubated for 20 min at 37°C, and fluorescence signals were detected using the SpectraMax MiniMax 300. Green fluorescence (Ex 495 nm; Em 530 nm) indicated live cells stained with Calcein AM, while red fluorescence (Ex 530 nm; Em 645 nm) indicated dead cells stained with PI. For further investigation, the assay was scaled up using 100-mm dishes under similar conditions. After 14 h of treatment with DIMATE or ABD0171, capillary electrophoresis immunoassay was performed as described above.

### Quantification of soluble IL-6 and tPA proteins

Cells were cultured in 100-mm dishes under anchored (uncoated) or non-anchored (poly-HEMA-coated) conditions. After adherence, cells were treated for 14h with DIMATE, ABD0171, or vehicle control. Supernatants were collected, centrifuged, and assayed for IL-6 and tPA protein levels using the Human IL-6 and Human t-Plasminogen Activator DuoSet ELISA Development Kit (R&D Systems, Minneapolis, MN, USA), following the manufacturer’s instructions.

### Animal experiments

All mice were obtained from Janvier Labs (Le Genest-Saint-Isle, France) or Charles River GmbH (Sulzfeld, Germany) and housed at the Commune Scar Rockefeller animal facility (Lyon, France) or Reaction Biology Europe GmbH (Freiburg, Germany), following institutional and governmental animal welfare regulations.

For toxicity studies, four female SCID-CB17 immunocompromised mice per group were treated with intra-peritoneal injections of DIMATE (15 or 20 mg/kg) or ABD0171 (30 or 40 mg/kg), t.i.w. for four weeks. The control group received vehicle (PBS containing 10% Kolliphor® HS15). Mice were monitored for body weight, physical condition, and blood sampling was conducted (days 5, 19, and 26) for haematological and enzymatic analyses.

For xenograft models, 5 × 10⁶ MDAMB231-GFP cells were orthotopically implanted into 50 SCID-CB17 immunocompromised mice. For syngeneic models, 0.5 × 10⁶ 4T1_Luc-M3 cells were orthotopically implanted into 36 BALB/cAnNCrl immunocompetent mice.

Once tumors reached ∼50 mm³, mice were randomized into treatment cohorts: DIMATE (20 mg/kg), ABD0171 (40 mg/kg), or vehicle control (n = 10 per group for MDAMB231-GFP, n = 12 per group for 4T1_Luc-M3). Treatments were administered intra-peritoneally, t.i.w. for three weeks. Tumor growth, body weight, and overall health were monitored daily. Sacrifice occurred at the study endpoint (week 3) or earlier if tumor volumes reached 700 mm³ (MDAMB231-GFP) or 1600 mm³ (4T1_Luc-M3). For the 4T1_Luc-M3 model, various organs were harvested for *ex vivo* luciferase assays to assess metastasis.

### Immunohistochemical stainings

Formalin-fixed, paraffin-embedded (FFPE) tumor samples were analyzed by immunohistochemistry (IHC) at CIQLE, Lyon (UAR3453 CNRS, US7 Inserm, UCBL). Hematoxylin and eosin (H&E) staining was performed on 3 µm sections using standard protocols. For IHC, sections were incubated with the following primary antibodies: anti-Ki67 (Abcam, Paris), anti-cleaved caspase-3 (Cell Signaling, Danvers, MA), anti-cleaved PARP (Cell Signaling), anti-CD4 (Abcam), and anti-CD8α (Cell Signaling). Secondary antibodies conjugated to biotin were applied, and staining was performed using the Roche Ventana Discovery XT immunostainer (Roche Diagnostics, Meylan). Antibody detection was achieved with the DAB Map Detection Kit (Roche Diagnostics). Image analysis and quantification were carried out using the open-source software QuPath.

### Ex vivo bioluminescence imaging (BLI) analysis

Organs from experimental animals were analyzed for bioluminescent signals. The control group data were separated to facilitate outlier detection and removal using the interquartile range (IQR) method. Median values from the control samples were calculated for each organ and plate, and these were used to normalize the replicate measurements of the experimental samples. Negative normalized values were adjusted to zero to ensure all values represented positive biological signals. To identify potential outliers, group-specific criteria were applied. For Group 1 (PBS), both lower and upper bounds for outlier detection were defined based on the IQR, while for Groups 2 and 3 (DIMATE and ABD0171), only upper-bound outliers were flagged. Subsequently, outliers were removed from each group. Ex vivo BLI values were calculated for each sample by adjusting the median of the normalized replicates according to sample volume (10 µL) and animal body weight (g), yielding values expressed as BLI per gram of tissue. Statistical comparisons between treatment groups and controls were performed using the Wilcoxon rank-sum test, with p-values adjusted for multiple comparisons using the Benjamini-Hochberg method. A threshold of p ≤ 0.05 was used to determine significance.

### Statistical analyses of experimental data

GraphPad Prism (GraphPad Software, Inc., La Jolla, CA, USA) was used for the graphic representations and statistical analysis of the data. Mann-Whitney test was used for the comparison between two groups and Kruskal-Wallis test for multiple comparisons. The differences between different groups with the control were considered as significant if the p-values were ⩽ 0.05.

## ACKNOWLEDGEMENTS

This research was partially funded by the Spanish Ministerio de Ciencia e Innovación (Agencia Estatal de Investigación, grant number PID2020-119424RB-I00 / AEI / 10.13039/501100011033), Plan France Relance by the French Ministère de l’Économie et des Finances, and received funding from Xerys Invest. The funders had no role in the study design, data collection and analysis, decision to publish, or preparation of the manuscript. The results published here are in part based upon data collected from the TCGA Research Network: https://www.cancer.gov/tcga.

## AUTHOR CONTRIBUTIONS

RRR carried out bioinformatics and statistical analyses. RP, XP, and JF designed, conducted, and interpreted enzymatic and biochemical assays. OP performed mass spectroscopy assays. AC conducted and analyzed the *in vitro* cytotoxicity screenings and functional assays. AC, PB, BJ and AK performed the *in vivo* studies and immunohistochemical (IHC) analyses. DC, AP, GF, and GM were responsible for the chemical synthesis and corresponding analytical studies. MP, RRR, RP, and AC interpreted data and prepared the figures. MP, JF, and RRR conceptualized, interpreted and supervised the study, with major contributions to the writing and editing of the manuscript.

