## Supplementary Information for "ALDH1 subtype-specific inhibitor targets key cancerous epithelial cell populations in aggressive subtypes of breast cancer"

### SUPPLEMENTARY MATERIAL AND METHODS

**ROS assay.** Intracellular ROS levels were assessed using the fluorescent probe 2',7'-dichlorodihydrofluorescein diacetate (H2DCFDA, VWR, FR), following the manufacturer's protocol. Briefly, cells were incubated with H2DCFDA at a final concentration of 20μM for 30 minutes at 37°C in the dark. Fluorescence intensity of the oxidized form (DCF), directly proportional to ROS levels, was measured using an Accuri C6 flow cytometer and analyzed using Kaluza software.

### SUPPLEMENTARY FIGURE LEGENDS

**Supplementary Figure 1. Expression of ALDH1A isoforms and dose-response curves of breast cancer cell lines treated with the ALDH inhibitors. (a)** WES capillary immunoelectrophoresis analysis of ALDH1A1, ALDH1A2 and ALDH1A3 protein expression levels across 18 human and one murine breast cancer cell lines, grouped by their corresponding intrinsic molecular subtype. **(b)** Dose-response curves for cell viability in the panel of 18 human breast cancer cell lines treated with increasing concentrations of ABD0171 or DIMATE (0 to 50 μM) for 48 hours. The growth-inhibitory effect was assessed using the Resazurin sodium salt reduction assay. Curves were fitted via non-linear regression analysis of log-dose/response curves using GraphPad software.

**Supplementary Figure 2. Kinetic analyses of DIMATE and ABD0171 as ALDH3A1 and ALDH2 inhibitors and mass spectrometry validation of DIMATE as a specific affinity label of ALDH1A3. (a)** Substrate titration of steady state velocity for ALDH3A1 in the presence of DIMATE and ABD0171 acting as non-competitive inhibitors at varying concentrations (left panels) and using 4-nitrobenzaldehyde as a substrate in the corresponding reaction buffer; and $k_{inact}/K_i$ values calculated from the plots of $k_{obs}$ *versus* concentrations of DIMATE and ABD0171 for ALDH2 (inner plots). Enzymatic activities were measured at saturating substrate concentrations (250 μM hexanal) in the corresponding reaction buffer. Data are presented as mean ± standard deviation (right panels). **(b)** $k_{inact}/K_i$ values calculated from the plots of $k_{obs}$ *versus* concentrations of DIMATE in a range between 0-50 μM for ALDH1A3 (inner plots). Enzymatic activities were measured at saturating substrate concentrations (250 μM hexanal) in the corresponding reaction buffer. Data are presented as mean ± standard deviation **(c)** ESI-MS spectra of ALDH1A3 treated with 50 μM DIMATE. Single-charged peaks (1+) were observed at m/z 61076–61107. Red signals represent ALDH1A3 treated with 50 μM DIMATE (at an enzyme concentration of 16.7 μM), while black signals correspond to the untreated ALDH1A3 control. The spectra confirm irreversible binding of ALDH1A3 to DIMATE. Samples were prepared in 100 mM ammonium bicarbonate (pH 7.5) and pre-incubated for 45 min at 37°C.

**Supplementary Figure 3. Effects of ALDH Inhibitors on oxidative stress, metastatic potential, and key oncogenic pathways in TNBC cells. (a)** Quantification of reactive oxygen species (ROS) levels in MDAMB231, MDAMB468 and HCC70 cells exposed to 15µM of DIMATE, ABD0171, or vehicle control for 2 hours. **(b)** Representative images from invasion and migration assays (top-left panels) in MDAMB468 and HCC70 cells treated with either ABD0171 or DIMATE for 24- 48h. Scatter plots (bottom-left) show quantification from two independent experiments (median ± standard deviation). Statistical analysis was performed using Kruskal-Wallis test. Evaluation of anoikis resistance in MDAMB468 and HCC70 cells cultured on anchored and non-anchored (poly-HEMA-coated) surfaces and treated with vehicle control, DIMATE or ABD0171 for 24h. Representative images of cells stained with calcein-AM and propidium iodide (top-right panels) illustrate cell viability and death. Quantification of cell death using the resazurin-based toxicity assay is presented as median values ± standard deviation from triplicate experiments (bottom-right panels). Statistical significance was determined using an unpaired t-test with Welch's correction. **(c)** WES capillary immunoelectrophoresis analysis of total and phosphorylated AKT and ERK protein expression levels in TNBC cells treated with DIMATE or ABD0171 for 14h, under anchored and non-anchored conditions.

**Supplementary Figure 4. Evaluation of body weight during in vivo toxicity and efficacy studies of ALDH inhibitors in breast cancer models. (a)** Body weight changes over time, expressed as percentage relative to day 0, in healthy SCID-CB17 mice during the *in vivo* toxicity study. One animal in the group treated with the highest dose of DIMATE (20mg/kg) showed a weight loss exceeding 10% and was therefore euthanized. **(b, c)** Body weight measurements during *in vivo* efficacy studies for DIMATE and ABD0171 in the MDAMB231 xenograft model and the immunocompetent 4T1_Luc-M3 mouse models of TNBC.

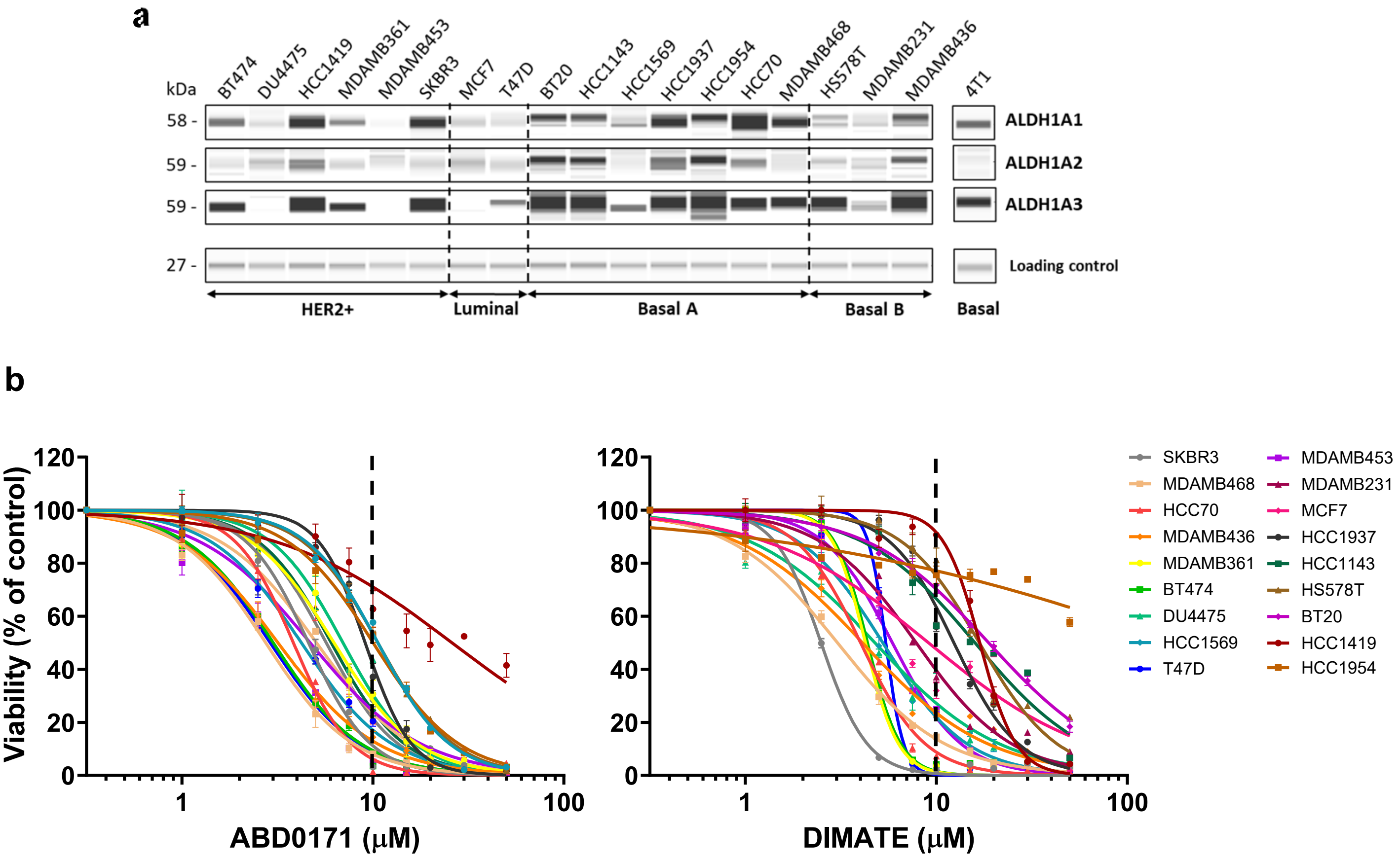


Supplementary Figure 1

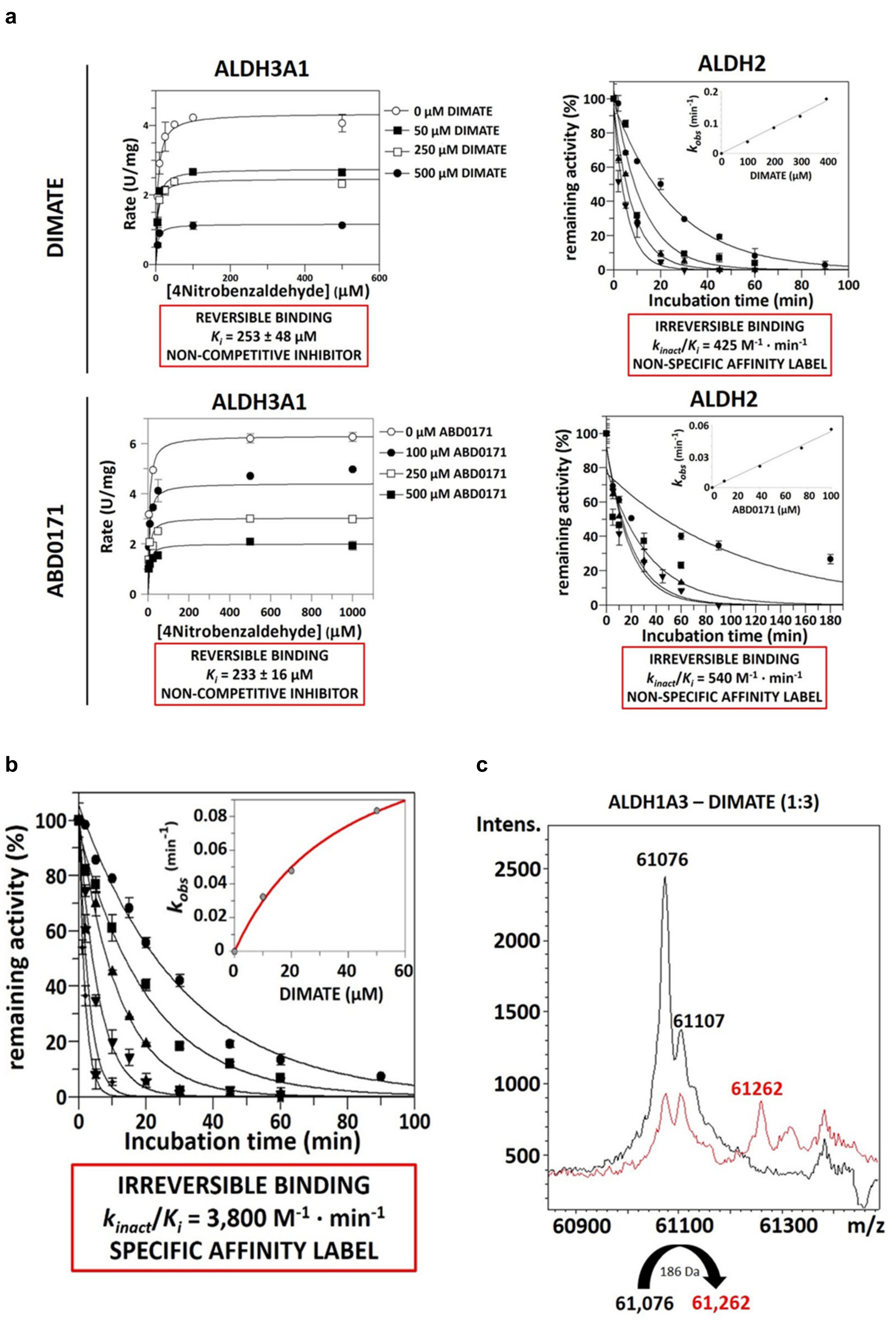


Supplementary Figure 2

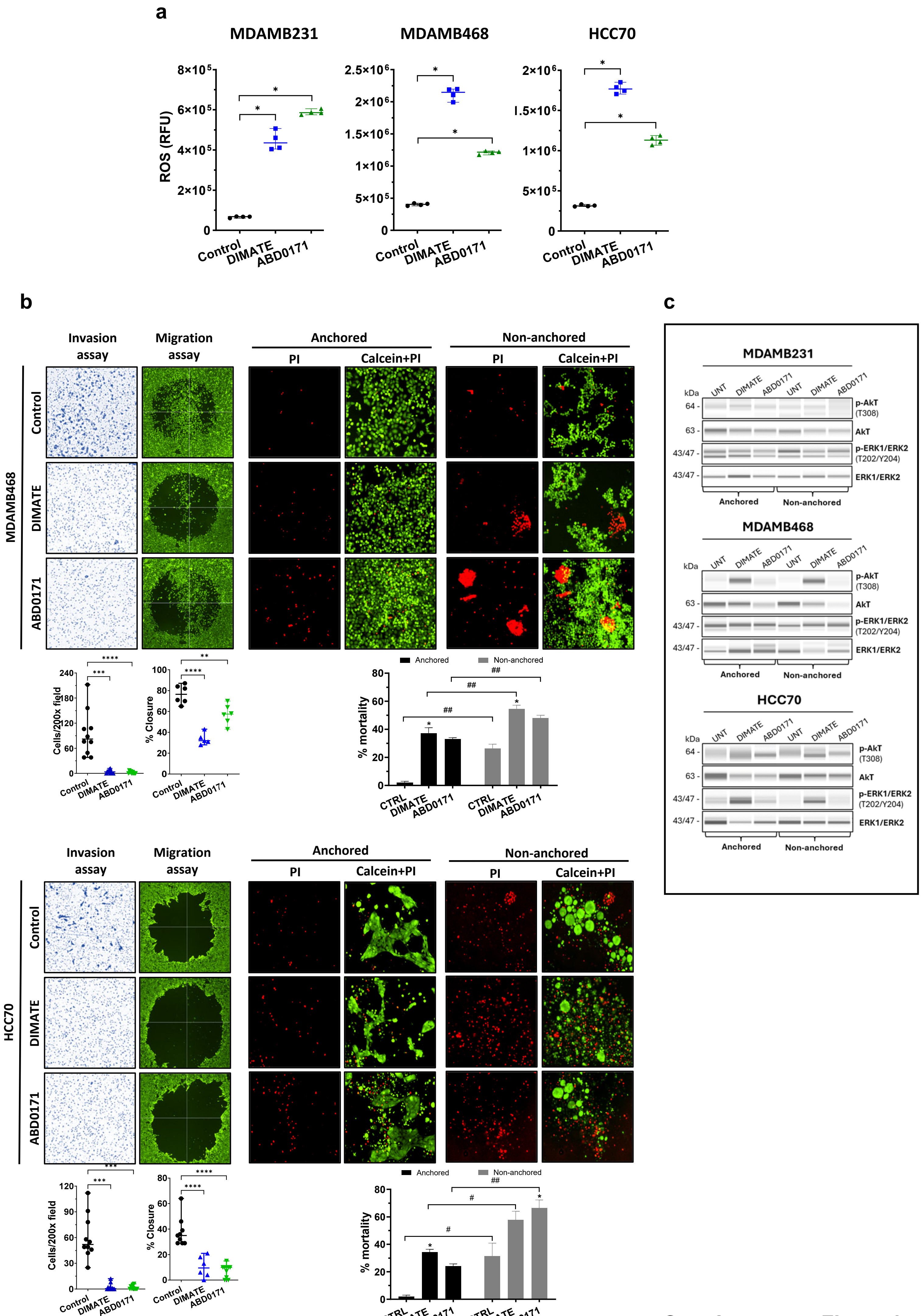


Supplementary Figure 3

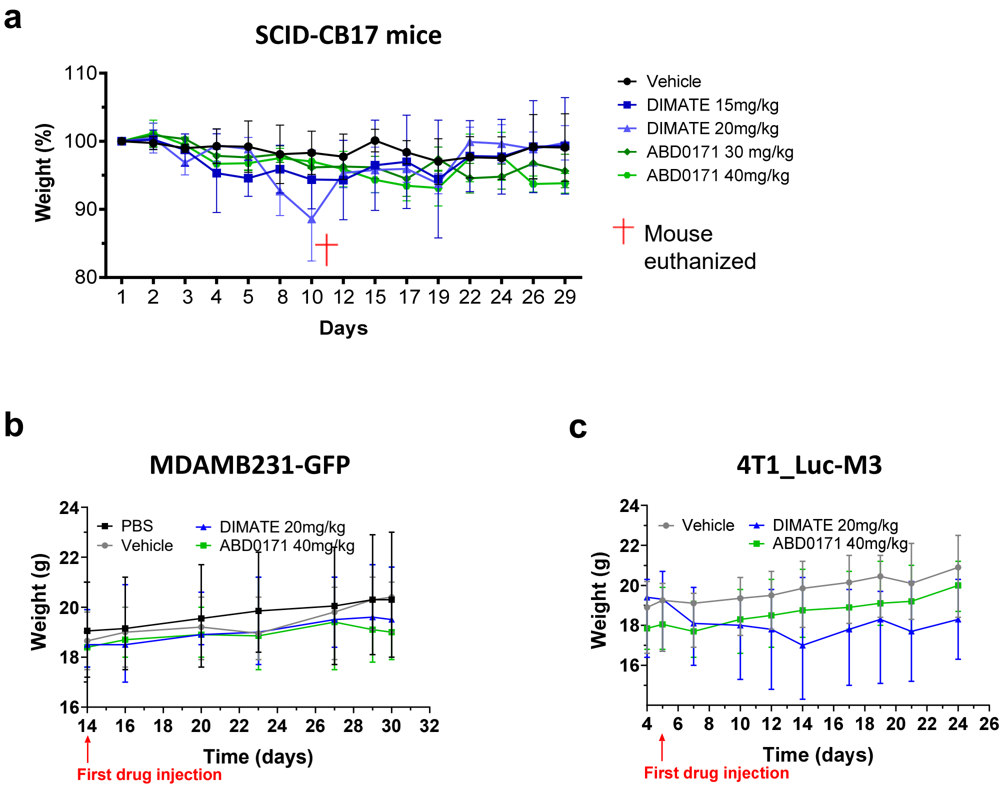


**Supplementary Figure 4**

**Table S1.** $IC_{50}$ (μM) and $k_{inact}/K_i$ ($M^{-1} \cdot min^{-1}$) or $K_i$ values of DIMATE and ABD0171 for ALDH3A1 and ALDH2 using 4-nitrobenzaldehyde and hexanal as substrate, respectively.

| | ALDH3A1 | | ALDH2 | |
|---|---|---|---|---|
| **COMPOUND** | **$IC_{50}$ (μM)** | **$K_i$ (μM)** | **$IC_{50}$ (μM)** | **$k_{inact}/K_i$ ($M^{-1} \cdot min^{-1}$)** |
| **DIMATE** | 303 ± 46 | 253 ± 48 | 72 ± 9 | 425 |
| **ABD0171** | 242 ± 11 | 233 ± 16 | 12 ± 3 | 540 |

$IC_{50}$ values were obtained fluorometrically using 4-nitrobenzaldehyde for ALDH3A1 and hexanal for ALDH2 as substrates. The reaction mixture was incubated in the corresponding reaction buffer[71] for 20 min at 37°C, before adding $NAD^+/NADP^+$ cofactor, NADH/NADPH internal standard and the corresponding substrate. $k_{inact}/K_i$ values were obtained by $k_{obs}$ *vs* [I] plot. $K_i$ values were obtained using a non-competitive fit of DIMATE and ABD0171 inhibition against ALDH3A1 at various concentrations of inhibitor compound. Experimental values are the mean ± standard error.

**Table S2.** List of antibodies used for protein detection in capillary electrophoresis immunoassays and/or in immunohistochemistry staining.

| Antibody | Provider | Reference |
|---|---|---|
| **ALDH1A1** | Proteintech | 15910-1-AP |
| **ALDH1A2** | Proteintech | 13951-1-AP |
| **ALDH1A3** | Novus | NBP2-15339 |
| **JNK** | Cell Signaling | 9252 |
| **Phospho-JNK (T183/Y185)** | Cell Signaling | 4668 |
| **p38** | Cell Signaling | 8690 |
| **Phospho-p38 (T180/Y182)** | Cell Signaling | 4511 |
| **c-Jun** | Cell Signaling | 9165 |
| **Phospho-c-Jun Ser73 (S73)** | Cell Signaling | 3270 |
| **Flip** | Cell Signaling | 56343 |
| **XIAP** | Cell Signaling | 14334 |
| **C-IAP2** | Cell Signaling | 3130 |
| **Cleaved Caspase 8** | Cell Signaling | 9496 |
| **Cleaved Caspase 3** (WES) | Cell Signaling | 9662 |
| **Cleaved Caspase 7** | Cell Signaling | 9492 |
| **Cleaved PARP** (WES/IHC) | Cell Signaling | 5625 |
| **STAT3** | Cell Signaling | 9139 |
| **Phospho-STAT3 (Y705)** | Cell Signaling | 9145 |
| **JAK2** | Cell Signaling | 3230 |
| **Phospho-JAK2 (Y1007/1008)** | Cell Signaling | 3776 |
| **FAK** | Cell Signaling | 3285 |
| **Phospho-FAK (Y397)** | Cell Signaling | 8556 |
| **Phospho-Src (Y416)** | Cell Signaling | 20101 |
| **Loading Control** | Protein Simple | 042-196 |
| **Ki67** | Abcam | ab15580 |
| **Cleaved Caspase-3** (IHC) | Cell Signaling | 9661 |
| **CD4 (T-cells)** | Abcam | ab183685 |
| **CD8 (T-cells)** | Cell Signaling | 98941 |
